# Effects of tree species diversity and identity on remotely sensed canopy variables in two forest biodiversity experiments

**DOI:** 10.64898/2026.11.09.765348

**Authors:** Sofia J. van Moorsel, Bernhard Schmid, Michael Niederberger, Tiziana L. Koch, Michael Scherer-Lorenzen, Daniel Kopkáně, Uwe Rascher, Alexander Damm, Meredith C. Schuman

## Abstract

- Remote sensing can link biodiversity and its effects on ecosystem functioning in forests, but in observational studies environmental covariates may confound results.
- We tested effects of tree species richness and functional diversity on remotely-sensed canopy variables at community and species levels in two forest biodiversity experiments established under homogeneous environmental conditions. At the BIOTREE sites in Kaltenborn and Bechstedt, we combined airborne imaging spectroscopy-based vegetation indices, sun-induced chlorophyll fluorescence, and land surface temperature at 1-m resolution, with ground-based tree stem growth as a productivity measure.
- At Kaltenborn, increasing species richness (one to four species) did not increase community productivity because of contrasting species-level responses. The normalized difference vegetation index (NDVI) decreased with richness in both needle-leaved species but increased in both broadleaved species. At Bechstedt, increasing functional diversity of mixtures planted with four tree species lowered community-level normalized difference water and photochemical reflectance indices; responses differed between four functional groups of tree species.
- NDVI was negatively correlated with basal area; correlations among other spectral, thermal, and fluorescence traits varied among tree functional groups.
- These results show that forest biodiversity experiments can test remotely sensed indicators of tree diversity and ecosystem functioning and inform interpretation of observational studies.

## 1 Introduction

Biodiversity–ecosystem functioning relationships in forests are gaining attention (Huang et al., 2018; Liu et al., 2026) with evidence suggesting that tree species and functional diversity enhance ecosystem functions such as productivity and nutrient cycling (Deng et al., 2025; Huang et al., 2017; Bongers et al., 2021). Understanding how tree species diversity influences forest functioning, and identifying the mechanisms driving biodiversity effects, is an important research priority (Deng et al., 2025; Grossiord et al., 2013; Schnabel et al., 2021).

Assessing biodiversity–ecosystem functioning relationships across forests requires spatially extensive, repeatable measurements. Ground-based assessments, such as measuring basal area, provide detailed information, but are labour-intensive and spatially limited. Remote sensing assessments, however, enable the observation of vegetation properties across broad spatial scales while maintaining consistent and repeatable measurements (Pettorelli et al., 2014). Airborne and satellite remote-sensing platforms can complement ground-based measurements in ecological studies (Cavender-Bares et al., 2025). Imaging spectroscopy in particular has emerged as a powerful approach for linking spectral signals with plant functional traits and ecosystem functioning (Ustin et al., 2004; Gamon et al., 2019; Cherif et al., 2023). Spectral data provide an integrated representation of canopy variables, including canopy architecture, leaf chemistry, and water status, and may capture variation in plant functional strategies and the outcomes of species interactions within communities (Williams et al., 2020; Cavender-Bares et al., 2017; Serbin and Townsend, 2020). Mapping this functional variation is important because differences in canopy traits influence ecosystem processes and can help explain how plant communities respond to environmental change (Aguirre-Gutiérrez et al., 2025). In tree diversity experiments and natural mixed-species forests, imaging spectroscopy offers the opportunity to quantify species- and community-level responses of remotely-sensed canopy variables to tree diversity and to detect shifts in species functional traits and community functional trait compositions associated with interspecific interactions (Urgoiti et al., 2023; Felix et al., 2023; Veryard et al., 2023; Williams et al., 2020). This can provide insight into the mechanisms underlying biodiversity–ecosystem functioning relationships (Forrester and Bauhus, 2016), for example by revealing how species differ in canopy physiology, resource use, or stress responses when growing in monocultures versus mixtures (Felix et al., 2023) or how canopy structural complexity mediates biodiversity–productivity relationships (Deng et al., 2025).

Spectral vegetation indices capture specific aspects of canopy physiology and structure (Bannari et al., 1995). Reflectance-based vegetation indices can provide information on canopy greenness (*e.g.*, normalized difference vegetation index (NDVI), enhanced vegetation index (EVI), modified soil adjusted vegetation index (MSAVI)), pigment information (*e.g.*, chlorophyll index red-edge (Clre), chlorophyll content index (CCI), photochemical reflectance index (PRI)), and water status (*e.g.*, normalized difference water index (NDWI)) (Table 1, (Pettorelli et al., 2005)). Together, these vegetation indices provide complementary proxies for structural and physiological dimensions of canopy functioning and offer the potential to assess how species richness, species composition, and species identity are reflected in forest canopy variables (Koch et al., 2025; Helfenstein et al., 2022). Sun-induced chlorophyll fluorescence (SIF) provides information related to photosynthetic activity (Rascher et al., 2015) and correlates with gross primary productivity across a wide range of terrestrial ecosystems (Paul-Limoges et al., 2018; Damm et al., 2010, 2015a). Beyond its relationship with productivity, SIF-based approaches show potential to help constrain the process of plant transpiration, although mechanistic underpinnings are still the subject of ongoing research(Ahmed et al., 2023; Damm et al., 2021). SIF can capture rapid physiological responses to environmental constraints, including reductions in photosynthetic functioning associated with limited water availability (Quiros-Vargas et al., 2026; Damm et al., 2022). Furthermore, combined measurements of SIF, reflectance, and canopy temperature have demonstrated the potential of airborne remote sensing to detect stress-induced changes in canopy photosynthesis (Pinto et al., 2020). Therefore, SIF provides complementary information to reflectance-based indices by directly integrating variation in photosynthetic activity and physiological stress responses (Rossini et al., 2015; Zeng et al., 2022). Canopy temperature assessed through land surface temperature (LST) derived from thermal measurements in the long-wave infrared provides an additional perspective by capturing the balance between absorbed radiation, sensible heat dissipation, and evaporative cooling, and is therefore closely linked to plant transpiration and water status (Javadian et al., 2022; Gräf et al., 2026).

**Table 1.** Overview of canopy variables and basal area. The ten canopy variables include seven vegetation indices derived from canopy reflectance data (NDVI, EVI, NDWI, CIre, CCI, MSAVI and PRI), land surface temperature (LST) and two sun-induced fluorescence (SIF) estimates. The ten canopy variables were supplemented with basal area as a proxy for biomass. For each vegetation index, the reflectance value of the spectral band closest to the target wavelength defined in the original literature was used (as in the shown formulae).

| **Canopy variable** | **Index** | **Abbreviation** | **Used formula** | **Original reference** |
|---|---|---|---|---|
| Canopy greenness | Normalized Difference Vegetation Index | NDVI | $\frac{R_{800}-R_{672}}{R_{800}+R_{672}}$ | Rouse et al. (1974); Tucker et al. (1979) |
| Canopy greenness | Enhanced Vegetation Index | EVI | $2.5\cdot\frac{R_{760}-R_{672}}{R_{760}+6R_{672}-7.5R_{470}+1}$ | Huete et al. (2002) |
| Canopy water content | Normalized Difference Water Index | NDWI | $\frac{R_{857}-R_{1240}}{R_{857}+R_{1240}}$ | Gao (1996) |
| Chlorophyll | Red-edge Chlorophyll Index | CI<sub>*re*</sub> | $\frac{R_{783}}{R_{704}}-1$ | Gitelson et al. (2003) |
| Chlorophyll/carotenoid ratio | Chlorophyll/Carotenoid Index | CCI | $\frac{R_{528}-R_{645}}{R_{528}+R_{645}}$ | Gamon et al. (2016) |
| Canopy greenness | Modified Soil Adjusted Vegetation Index | MSAVI | $\frac{2R_{857}+1-\sqrt{(2R_{857}+1)^2-8(R_{857}-R_{672})}}{2}$ | Qi et al. (1994) |
| Canopy photosynthetic stress | Photochemical Reflectance Index | PRI | $\frac{R_{530}-R_{571}}{R_{530}+R_{571}}$ | Gamon et al. (1992) |
| Photosynthetic activity | Sun-induced fluorescence | SIF687 | – | – |
| Photosynthetic activity | Sun-induced fluorescence | SIF760 | – | – |
| Canopy temperature | Land surface temperature (K) | LST | – | – |
| Biomass/stand structure | Basal area (m<sup>2</sup> ha<sup>−1</sup>) | BA | – | – |

These vegetation indices and solar induced fluorescence have largely been established and tested in observational studies, in which environmental changes may be correlated to targets of measurement, such as canopy moisture or photosynthetic activity. Here, we test these metrics in experimental forests where either species diversity or functional diversity has been manipulated for plots of trees grown in a common environment. In such experiments, biodiversity and particularly species diversity is usually observed to increase productivity. We wished to know whether commonly used indices reveal species differences and biodiversity effects when these have been experimentally manipulated and thus uncorrelated from other environmental variation. Furthermore, we asked whether remotely sensed variables such as NDVI and other greenness indices are good predictors of measured productivity in an experimental setting with reduced environmental correlations.

We used airborne remote sensing data from two of the worldwide oldest tree diversity experiments, situated in temperate Europe. We tested whether remotely sensed canopy variables vary with tree species richness or with functional diversity within a given species richness level. The Kaltenborn experiment with a species pool of four species represents a gradient in species richness from 1–4, whereas the Bechstedt experiment with a species pool of 16 species represents a functional diversity gradient among four-species mixtures of different species compositions (Scherer-Lorenzen et al., 2007). Combining airborne imaging spectroscopy, SIF and thermal infrared measurements we assessed whether remotely-sensed canopy variables reflect biodiversity-driven variation in ecosystem functioning and how this was related to species-specific responses to tree species diversity. The assessment of species-specific responses was possible due to the unique design of the experiment in which small groups of trees belonging to the same species were planted in patches whose “quilt” then formed the communities varying in species richness or functional diversity. We hypothesized that (i) tree species richness or functional diversity would alter remotely sensed canopy signals at community level and (ii) that species identity together with tree species richness or functional diversity would alter canopy signals at species level, explaining the responses at canopy level. Furthermore, we hypothesized that (iii) reflectance-based vegetation indices would be related to basal area. Finally, because reflectance-based vegetation indices, SIF and LST capture complementary but physiologically interconnected dimensions of canopy functioning, we hypothesized that (iv) correlations among these variables would differ among tree functional groups, reflecting differences in their leaf traits, canopy architecture, and physiological functioning.

## 2 Materials and methods

### 2.1 Study sites

This study was conducted at two BIOTREE forest biodiversity experiments established in 2003/2004 in Thuringia, Germany (Scherer-Lorenzen et al., 2007). The two experiments were specifically designed to disentangle different components of tree species diversity effects on ecosystem functioning. In the Kaltenborn experiment, tree species richness was manipulated using four tree species, planted as monocultures and as mixtures containing two, three, or four species. Thus, Kaltenborn represents a species richness gradient (1–4 species) in which richness levels are represented by different species compositions. In contrast, in the Bechstedt, functional diversity was manipulated using 16 species planted in different combinations of four species. Together, these two experiments allowed us to assess how species richness and functional diversity influenced remotely-sensed measures of canopy functioning at the level of species populations within a community and at the level of the entire community.

At the Kaltenborn site (50.77867, 10.22259, WGS84), 16 experimental plots are divided into three subplots representing different forest management treatments: unmanaged (u), conventionally managed (m; thinned according to regional close-to-nature forestry practices), and managed with additional low-abundance species (m+; Fig. 1a; Scherer-Lorenzen et al., 2007). We included only the unmanaged (u) and conventionally managed (m) treatments and used this subplot identity as random factor in models (see statistical analyses). We did not investigate the influence of the management because the management treatment (thinning) only began in winter 2020–2021 (Matevski et al., 2026). Each subplot consisted of 30 monospecific tree patches of 8 *→* 8 m, which were combined in a checkerboard design to create communities at subplot scale containing one to four tree species (Fig. 1a, 960 patches in total). The four species are the needle-leaved tree species Douglas fir (*Pseudotsuga menziesii*) and Norway spruce (*Picea abies*) and the broadleaved tree species European beech (*Fagus sylvatica*) and sessile oak (*Quercus petraea*). Each community composition was represented by one plot (four monocultures, six two-species mixtures, four three-species mixtures), except for the four-species mixture, which was represented by two plots.

**Fig. 1.**
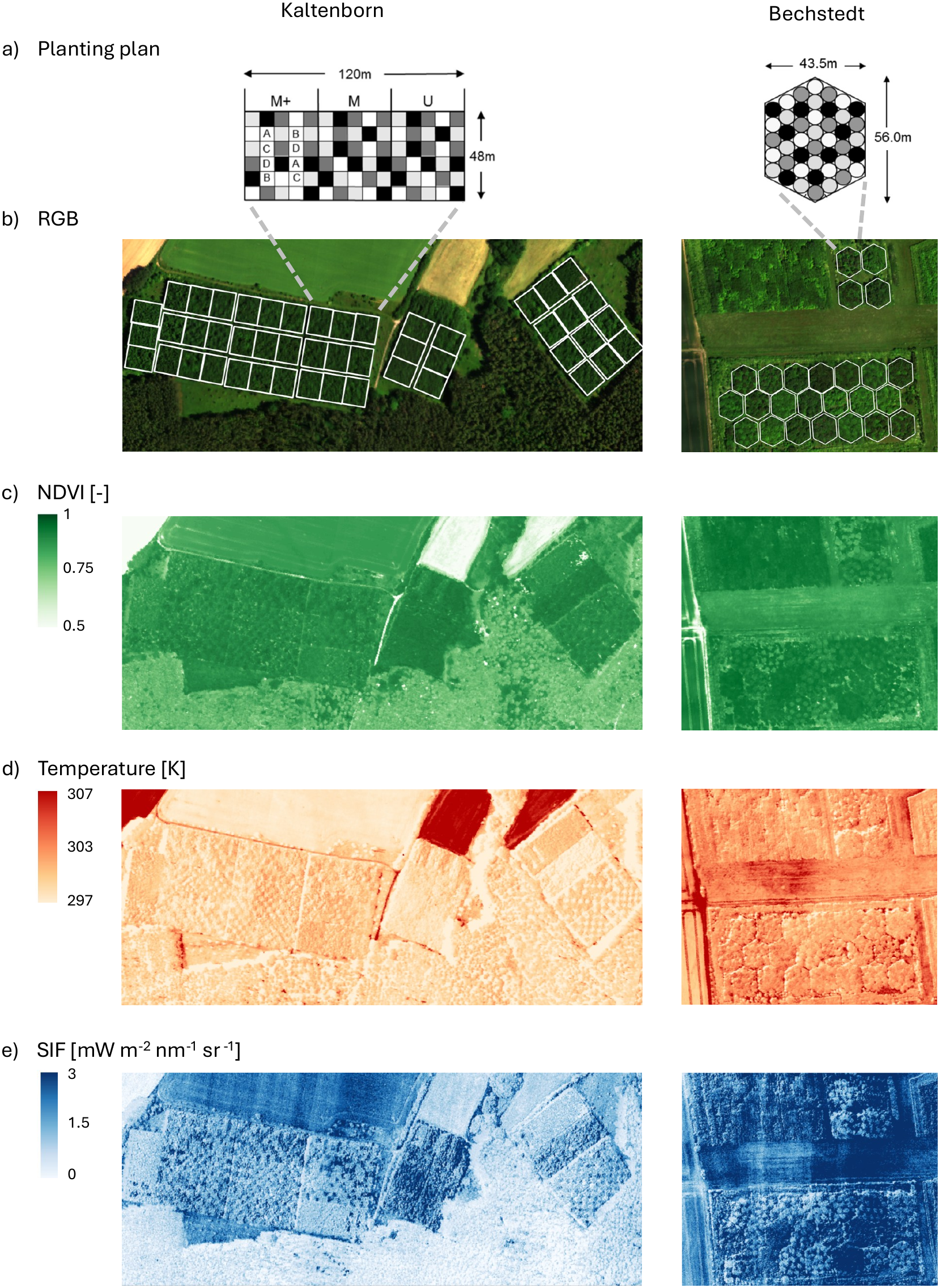
Overview of the Kaltenborn and Bechstedt field sites. a) Planting plan for both experiments. b) True color (RGB) image from HyPlant acquisition in 2021. c) Normalized difference vegetation index (NDVI) based on HyPlant-DUAL sensor reflectance data. d) Land surface temperature (canopy temperature) in K measured by the TASI sensor. e) Sun-induced fluorescence (SIF) as measured by the FLUO sensor. M = Managed, U = Unmanaged, M+ = Managed with additional species added. See text for sensor details.

At the Bechstedt site (50.8971, 11.08238, WGS84), 25 hexagonal plots were all planted with four tree species each, but representing different levels of functional diversity (Fig. 1a). Each plot consisted of 44 monospecific circular tree patches of 3.5 m radius, which were combined in a hexagonal pattern to create communities at plot scale (Fig. 1a, 1100 patches in total). The species pool consisted of 16 tree species: *Acer campestre*, *Acer platanoides*, *Acer pseudoplatanus*, *Betula pendula*, *Carpinus betulus*, *Fagus sylvatica*, *Fraxinus excelsior*, *Larix decidua*, *Pinus sylvestris*, *Populus tremula*, *Prunus avium*, *Quercus petraea*, *Sorbus aucuparia*, *Sorbus torminalis*, *Tilia cordata*, and *Ulmus glabra*. Functional diversity was manipulated by selecting four-species combinations spanning a gradient from functionally similar to intermediate and highly dissimilar species. First, functional diversity was calculated for all possible four-species combinations (1820 combinations) based on a dendrogram-derived index (Petchey and Gaston, 2002). Then, twenty-four combinations were selected to represent the gradient as uniformly as possible, with one combination planted twice. Species were classified according to functional similarity using nine above- and belowground traits related to productivity, resource use, and nutrient cycling, including leaf type, light requirement, mean annual growth increment, leaf nitrogen concentration, and litter C/N ratio (Scherer-Lorenzen et al., 2007). For analyses involving broader species groupings, the 16 species were grouped according to their position in this functional trait dendrogram. The resulting four groups were descriptively labelled as (1) small and slow-growing broadleaved species (*A. campestre*, *C. betulus*, *S. aucuparia*, and *S. torminalis*), (2) early-successional broadleaved species (*B. pendula*, *F. excelsior*, *P. tremula*, and *P. avium*), (3) late-successional broadleaved species (*A. platanoides*, *A. pseudoplatanus*, *F. sylvatica*, *Q. petraea*, *T. cordata*, and *U. glabra*), and (4) needle-leaved species (*L. decidua* and *P. sylvestris*). These labels reflect the functional clustering rather than predefined ecological strategy categories. Further details regarding experimental design, planting procedures, and site characteristics are provided by Scherer-Lorenzen et al. (2007).

### 2.2 Airborne data acquisition and preprocessing of remote sensing data

Airborne remote sensing data were acquired on 17 June 2021 during a single flight campaign over the Kaltenborn and Bechstedt sites (see also van Moorsel et al., 2026). The aircraft was equipped with the HyPlant imaging spectrometer, consisting of the DUAL and FLUO sensor modules, as well as a TASI thermal infrared sensor (Fig. 1bcd) (Siegmann et al., 2019). Meteorological conditions measured by the weather station on site during the campaign were warm and dry, with air temperatures exceeding 28 °C by 10:00 h and reaching a maximum of 30.8 °C at approximately 13:00 h, before remaining around 28 °C at 18:00 h. No precipitation was recorded at the local weather station during the seven days preceding the flight.

The HyPlant DUAL module records reflected radiance in 626 contiguous spectral bands covering 374–2504 nm. The spectral resolution is approximately 3.6 nm in the visible and near-infrared regions and 10.5 nm in the shortwave infrared region (Siegmann et al., 2019). The FLUO module records radiance between 670 and 780 nm with a spectral resolution of 0.28 nm, enabling retrieval of SIF within the oxygen absorption features around 687 nm (O_2_-B) and 760 nm (O_2_-A) (Siegmann et al., 2019). The TASI-600 sensor (ITRES Research Ltd., Calgary, Canada) measures thermal infrared radiation in the long-wave infrared region (22 bands from 8.6–10.9 *µ*m; Pignatti et al., 2011).

Four flight lines were combined into a mosaic for the Kaltenborn site and two flight lines were combined into a mosaic for the Bechstedt site. The average flight altitude was 680 m above ground level, resulting in a ground sampling distance of approximately 1 m for the DUAL and FLUO sensors and 0.8 m for the TASI sensor. Different acquisition times of individual flight lines during the campaign caused substantial differences in the thermal imagery between overlapping flight lines. We therefore radiometrically corrected individual TASI flight lines prior to mosaicking. In detail, the minimum number of TASI flight lines needed for full coverage was used (three at Kaltenborn, two at Bechstedt). For these lines, a homogeneous grass-covered rectangular area, visible in all flight lines at each site, was used to estimate a mean reference temperature per flight line. One flight line per site was designated as the reference and all other lines were corrected by an additive offset equal to the difference between their ROI mean and the reference ROI mean. Applied flight line temperature corrections were around ±0.5 K.

The HyPlant data were processed following established preprocessing workflows described by Siegmann et al. (2019) and van Moorsel et al. (2026). Briefly, raw DUAL data were radiometrically and geometrically corrected and converted from at-sensor radiance to top-of-canopy reflectance using the ATCOR-4 atmospheric correction approach (Richter and Schläpfer, 2002). The resulting reflectance data covered the 400–2500 nm spectral range. Spectral regions strongly affected by atmospheric absorption were interpolated during preprocessing. Only vegetation indices based on measured spectral bands were used in subsequent analyses.

The FLUO data were similarly processed according to the workflow described for the DUAL data to retrieve calibrates at-sensor radiances. SIF was retrieved using the improved Fraunhofer Line Discrimination (iFLD) method originally developed by Alonso et al. (2008) and adapted to airborne data as described in (Damm et al., 2015b). SIF retrievals were obtained for the O_2_-B and O_2_-A absorption features at approximately 687 and 760 nm, respectively. Additional details regarding HyPlant FLUO processing are provided by Siegmann et al. (2019). Thermal imagery from the TASI sensor was processed to land surface (kinetic) temperature (LST) using standard radiometric, geometric, and atmospheric correction and temperature–emissivity separation procedures as described by Hanuš et al. (2023).

At both sites, multiple individuals of the same species were grown in close proximity with the canopy cover largely closed. This made it unnecessary to delineate individual tree crowns and we could obtain single-species data at 1 m-resolution. The geolocation of these monospecific patches was determined based on the experimental planting design. For both sites, patch locations were reconstructed from planting plans and aligned with the airborne imagery using high-resolution satellite imagery and visual refinement of plot positions and orientations. At Kaltenborn, the centre locations of the 8 *→* 8 m patches were calculated within each subplot. At Bechstedt, center locations of the patches were derived from the hexagonal plot layout and the circular planting geometry. Spatial resolution of airborne imagery may result in mixed pixels at patch boundaries and small geolocation uncertainties can occur during image alignment. Therefore, we used a conservative pixel selection approach and we kept only the nine most central pixels of each patch for analyses to reduce potential edge effects and to make sure to only capture the target species within the patches. Then, to exclude shaded canopy elements that are often affected by errors in retrieved vegetation information (Damm et al., 2015b), the three brightest pixels among the nine central pixels were selected based on the summed reflectance between 400 and 2500 nm following Schneider et al. (2017) and van Moorsel et al. (2026) and used for all analyses presented here. The selected pixel locations were then matched between the spectral, SIF, and LST datasets, ensuring that identical canopy locations were used for all remotely sensed variables. Thus, SIF and LST values were extracted only from pixels identified as sunlit based on the reflectance data.

For reflectance-based analyses, two *P. abies* patches were identified as outliers and were excluded from subsequent analyses (see van Moorsel et al., 2026). The outliers corresponded to two *P. abies* patches in Kaltenborn plot 13 (*P. abies* monoculture), with dead spruce trees due to a bark beetle attack in 2018 (Jactel et al., 2025).

### 2.3 Canopy variables and basal area

Remotely-sensed canopy variables were selected to characterize complementary dimensions of canopy functioning, including structural and biochemical properties derived from reflectance, photosynthetic activity represented by SIF, and canopy energy balance represented by LST (Table 1). We calculated seven vegetation indices from the hyperspectral reflectance data (Table 1). The NDVI (Rouse et al., 1973; Tucker, 1979), and the MSAVI (Qi et al., 1994), which reduces soil background effects in vegetation reflectance measurements, were included as indicators of vegetation greenness (Pettorelli et al., 2005). The EVI originally introduced by Huete et al. (2002) and here calculated using the narrow-band hyperspectral formulation proposed by Rascher et al. (2015), was included because it is less affected by saturation effects in dense vegetation. The CI_re_ was used as an indicator of canopy chlorophyll content (Gitelson et al., 2003). In addition to these four “greenness indices”, canopy water status was characterized using the NDWI (Gao, 1996). The CCI was included as a proxy for pigment composition and changes in the relative abundance of chlorophyll and carotenoids (Gamon et al., 2016). Finally, we calculated the PRI, which is related to xanthophyll cycle activity and photosynthetic light-use efficiency (Gamon et al., 1992). For all vegetation indices, the reflectance value of the spectral band closest to the target wavelength was used.

HyPlant further allows retrieval of SIF, a more direct indicator of photosynthetic activity. SIF originates from the re-emission of absorbed light energy by chlorophyll molecules during the photosynthetic process. We used SIF retrievals from both the O_2_-A and O_2_-B absorption features (SIF.760 and SIF.687, respectively). LST was used to characterize canopy thermal conditions. LST integrates variation in canopy energy balance and is influenced by processes such as transpiration and water availability (cf. Zhou et al. (2023) for a summary of leaf related effects on temperature). LST provides complementary information on canopy functioning alongside reflectance-based vegetation indices and SIF.

Finally, we used tree basal area as a ground-based measure of productivity (Dietrich et al., 2023; Huang et al., 2018; Quaglino et al., 2026). Aboveground tree growth and productivity are commonly assessed using basal area or basal area increment, derived from repeated measurements of diameter at breast height (DBH) (Dietrich et al., 2023; Huang et al., 2018, 2022). In tree diversity experiments with a common planting date, cumulative basal area provides a proxy of tree above-ground productivity over the period of stand development. DBH (at 1.3 meters) measurements were collected during field surveys in winter 2019 at both the Kaltenborn and Bechstedt sites, representing the closest available survey year to the airborne data acquisition given that field surveys were not conducted annually. Individual tree basal area was calculated from DBH measurements and the area summed within patches. Patch-level basal area was subsequently converted to a hectare basis (m^2^ ha*^→^*^1^) by accounting for the respective patch area. Because DBH measurements were not available for all individuals and patches, basal area estimates were calculated only for patches with available measurements.

### 2.4 Statistical analyses

To evaluate how tree species richness, functional diversity and identity affected remotely sensed measures of canopy functioning at community and species level, we fitted linear mixed-effects models using the `lme()` function from the `nlme` package (v3.1.168) (Pinheiro et al., 2026) in R (v4.5.3, R Core Team 2026). All remotely sensed variables (vegetation indices, SIF, and LST) and basal area were analysed separately as response variables. SIF was log-transformed to fulfil model assumptions of homoscedasticity and normally distributed residuals. Model predictions and visualization of treatment effects were generated using the fitted mixed-effects models. Predictor variables were selected according to the experimental design of each site. For Kaltenborn, models included species richness, species identity, and their interactions, whereas for Bechstedt, models included functional diversity, species identity, and their interactions. Plot and subplot identity were included as random effects to account for the hierarchical experimental design. For Kaltenborn, species richness was included both as a categorical factor and as a log-transformed linear effect to distinguish discrete treatment effects from continuous richness responses. Because deviations from the log-linear effect were small, only analyses with the log-linear variable (log SR) are presented here. Phylogenetic class effects (CL) were assessed using contrasts between the needle-leaved tree species *vs*. broadleaved tree species and species identity effects using further contrasts within the classes (FA = contrast between *F. sylvatica* and *Q. petraea*, PI = contrast between *P. abies* and *P. menziesii*). Plot (PL) and subplot (TR, referring to management treatment) were included as nested random terms. The final model was coded as

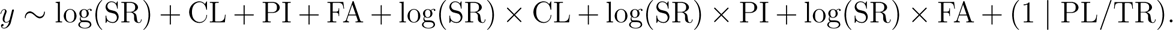

Here, the cross operator indicates interactions, the slash indicates nesting, and the vertical bar is the R-specific notation identifying random terms, *i.e.*, random intercepts for PL and TR within PL (= subplot identity). Note that the term log(SR) indicates richness effects at the community level, whereas all other terms indicate effects at species (patch) level, namely mean differences between taxa and differences in the responses of the different taxa to log(SR). As a consequence, significance tests for log(SR) effects at community level are based on comparing 16 plots whereas significance tests for species identity effects and log(SR) x species identity interactions at species level are based on comparing 958 patches. Thus, the statistical power to detect species-level effects is much higher than that to detect community-level effects. For the Bechstedt experiment, each spectral trait was analysed using a linear mixed-effects model of the form

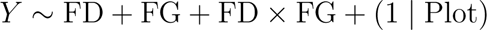

where FD refers to plot functional diversity and FG to functional group identity (n=4: small broadleaved tree species, early-successional broadleaved tree species, late-successional broadleaved tree species, and needle-leaved tree species). In Bechstedt, the design has no subplots, so plot identity was the only random term. Note that the term FD indicates functional diversity at community level, whereas the FG and the FD x FG terms indicate mean differences between functional groups and differences in the responses of the different functional groups to FD, respectively, at patch level. As for the Kaltenborn experiment, significance tests for FD effects at community level are based on comparing plots (here 25) whereas significance tests for FG effects and FD x FG interactions at species level are based on comparing patches (here 1100). Thus, the statistical power to detect species-level effects is again much higher than that to detect community-level effects.

### 2.5 Correlations between canopy variables and principal component analysis

Pairwise Pearson correlation coefficients were calculated among canopy variables using pairwise complete observations. Correlations were calculated separately for broadleaved tree species and needle-leaved species at Kaltenborn, and for the four functional groups at Bechstedt. P-values were obtained using the rcorr function from the Hmisc package (v5.2-5, Jr et al., 2026), and correlation matrices were visualized using the corrplot package (v0.95, Wei et al., 2024).

To summarize multivariate differences among remotely sensed canopy variables, principal component analyses (PCA) were conducted separately for the two experiments using ten variables derived from remote sensing. PCA was calculated using the prcomp() function in R. Variables were standardized using z-transformation prior to PCA. At Kaltenborn, PCA scores were visualized according to tree class whereas at Bechstedt, scores were visualized according to the four functional groups derived from the functional trait dendrogram. Variable loadings were plotted together with the PCA scores to visualize the contribution of individual canopy variables to the principal components.

## 3 Results

### 3.1 Effects of tree species richness and functional diversity on canopy variables at community level

Contrary to expectations, in Kaltenborn, species richness did not significantly affect any remotely sensed canopy variable at community level (Fig.2; Table S1). However, functional diversity at Bechstedt had significantly positive effects on community-level NDWI (*F* = 5.82, *P* = 0.024) and PRI (*F* = 5.63, *P* = 0.026). No significant effects were detected for the other canopy variables or basal area (Fig.3; Table S2).

**Fig. 2.**
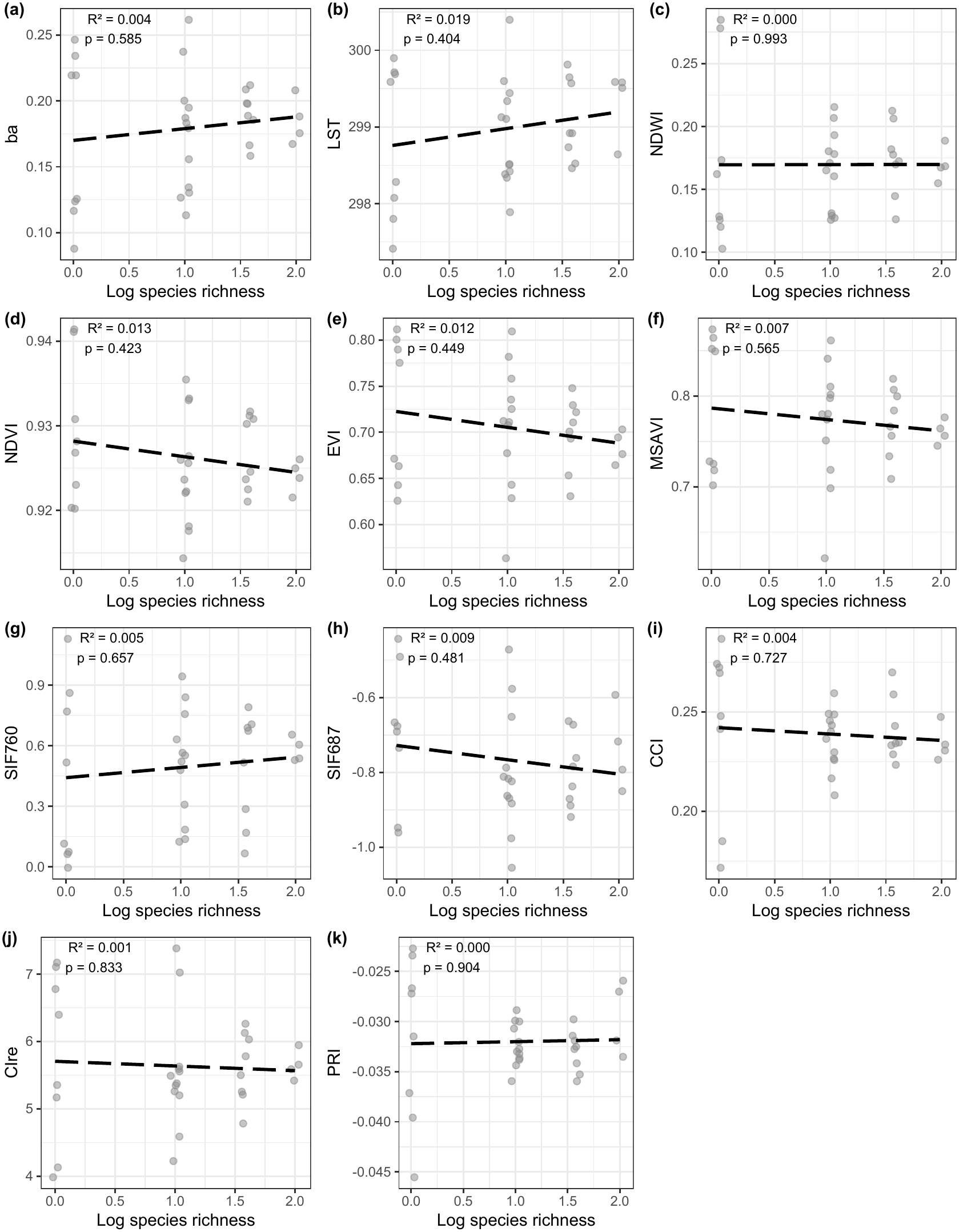
Overall effects of log2-transformed tree species richness on community-level means of canopy variables in the Kaltenborn experiment. The points show model-predicted mean responses of the 16 plots with each two subplots (n=32) in the Kaltenborn experiment. All non-significant relationships are shown with dashed lines. Marginal R2 for significant effects of tree species richness are reported in the corresponding panels. P-values for the fixed effect of logsr from a linear model including the random-effects term “plot/subplot” are shown in the panels and in Table S1.

**Fig. 3.**
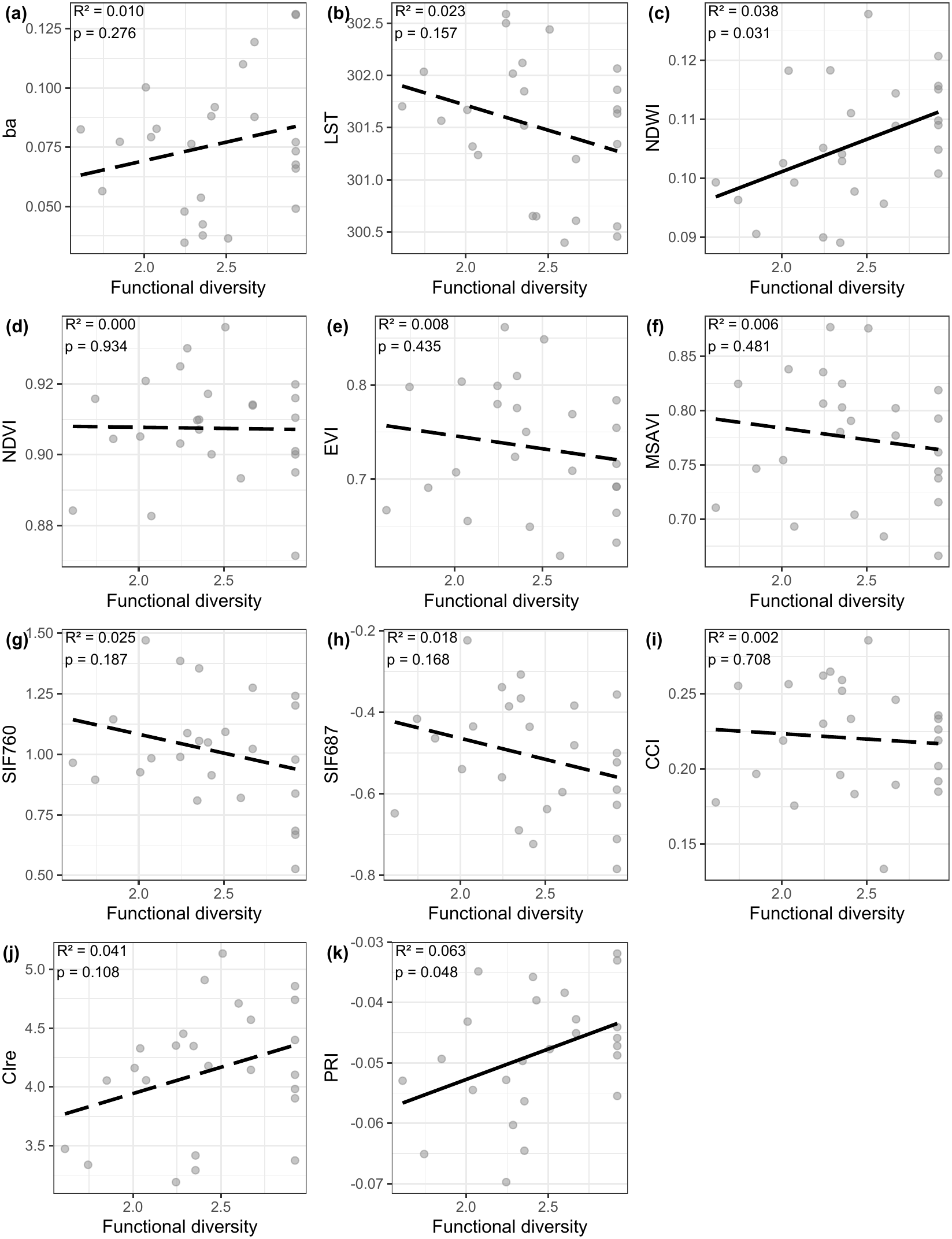
Overall effects of tree functional diversity on community-level means of canopy variables in the Bechstedt experiment. Points show model-predicted mean responses per plot (n=25). Significant relationships are plotted with solid lines, non-significant relationships with dashed lines. P-values for the fixed effect of functional diversity from a linear model are shown in the panels and in Table S2. Marginal R2 for significant effects of tree functional diversity are reported in the corresponding panels.

### 3.2 Effects of tree species richness and functional diversity on canopy variables at species level

At both sites, species differed in canopy characteristics and these were additionally modified by the biodiversity treatments (richness-dependent responses indicated by significant biodiversity x species identity interactions). In the Kaltenborn experiment, the four species strongly differed between and within classes (TableS1 and Fig.4a). The two needle-leaved tree species differed from the two broadleaved ones in all canopy variables except CCI. The effects were particularly pronounced for NDWI, NDVI, and basal area, indicating substantial differences in spectral properties and basal area. Within needle-leaved tree species and broadleaved tree species, species identity also had effects on canopy variables and basal area, significantly explaining variation in eight of the eleven canopy variables. Species identity effects were particularly strong for the vegetation indices, including NDWI, CCI, CIre, and PRI. Richness-dependent responses further differed between and within classes: basal area increased with richness in needle-leaved tree species but decreased in broadleaved tree species, whereas NDVI decreased with richness in needle-leaved tree species but remained relatively constant in broadleaved tree species (Fig.4a,d, Fig. S1a). LST increased with richness in *P. menziesii* but changed little in the other species, while CIre increased with richness in *Q. petraea* and decreased in *P. menziesii* (Fig.4b,j). Together, these contrasting responses indicate that effects of species richness on canopy variables depended strongly on the community, with larger differences between needle-leaved and broadleaved tree species (Fig.S1a,b) but also significant differences between species within these classes.

**Fig. 4.**
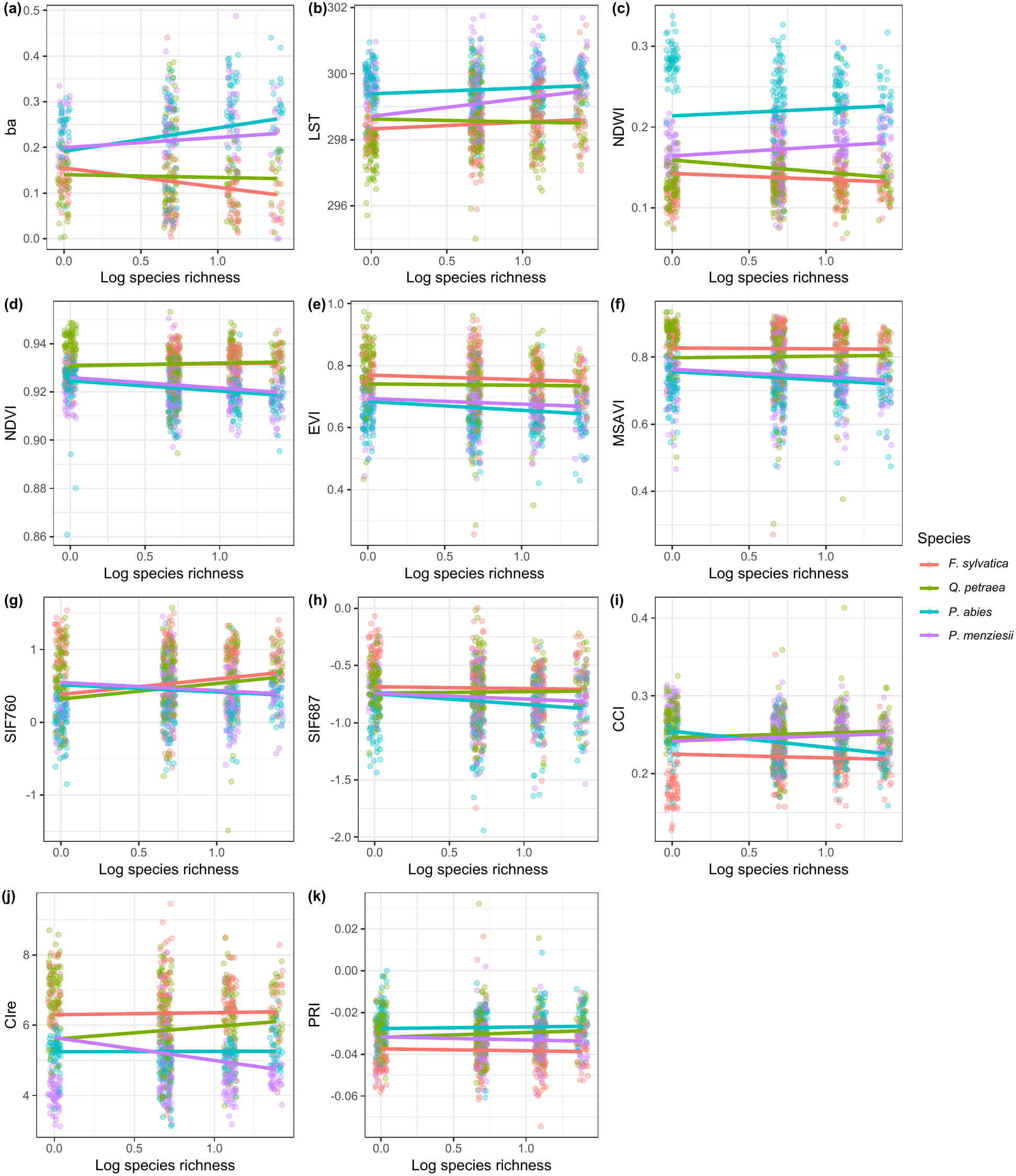
Species-level responses to species richness in the Kaltenborn experiment. Observed data and fitted mixed-effects model predictions showing the relationship between log-transformed species richness and canopy variables for each species. Lines indicate predicted values from the linear mixed models as described in the text, points represent individual patches (see also Table S1).

In the Bechstedt experiment, different plant functional groups (FG) significantly differed among each other in basal area and all canopy variables (Table S2, Fig. S1b). Furthermore, as in the Kaltenborn experiment, functional groups also showed different responses to the functional diversity gradient (significant functional diversity ! functional group identity interactions in Table S2, Fig. 5). Positive effects of functional diversity were most common in early-successional broadleaved tree species and, for several canopy variables, in needle-leaved tree species, whereas small broadleaved tree species generally showed weak or negative responses. The three greenness vegetation indices NDVI, EVI, and MSAVI increased with functional diversity in needle-leaved tree species, but remained relatively constant in the other three functional groups (Fig. 5d-f). Diversity responses of SIF variables were negative in small and late-successional broadleaved tree species, but relatively neutral in needle-leaved tree species and early-successional broadleaved tree species (Fig. 3g-h). The latter two groups instead showed positive diversity responses for CCI (Fig. 5i). Late-successional broadleaved tree species showed the most positive diversity responses for CIre and PRI (Fig. 5j-k).

**Fig. 5.**
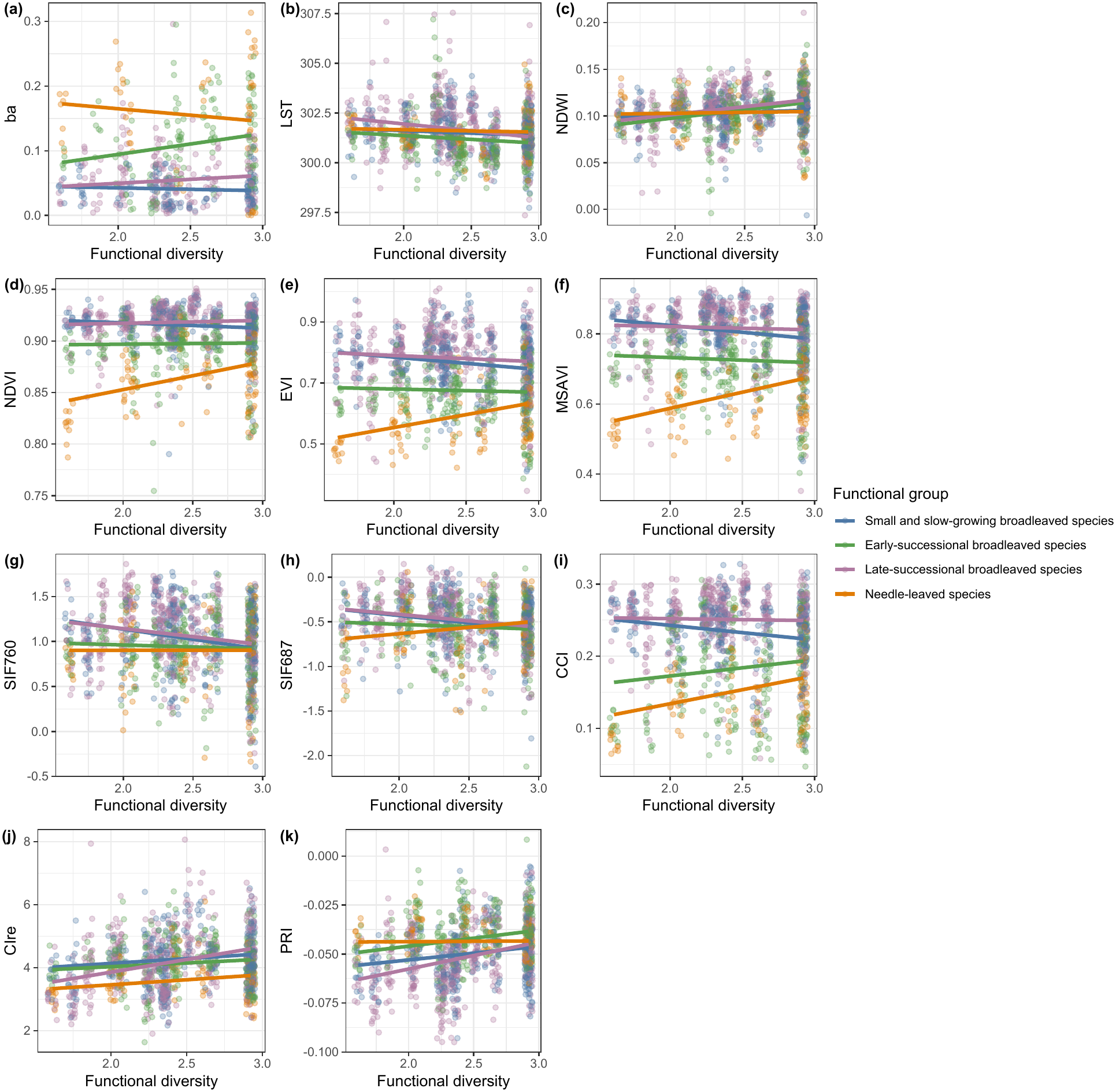
Functional group-level responses to functional diversity in the Bechstedt experiment. Observed data and fitted mixed-effects model predictions showing the relationship between functional diversity and canopy variables for each functional group. Lines indicate predicted values from the linear mixed models as described in the text, points represent individual patches (see also Table S2).

### 3.3 Correlations between canopy variables

In both experiments, the strongest and most consistent correlations were observed among the optical vegetation indices, whereas SIF and LST showed generally weak relationships with the optical indices (Fig. S2, Fig. S3).

In Kaltenborn, NDVI, EVI, MSAVI, and CIre were all positively correlated, with particularly strong relationships between EVI and MSAVI (r = 0.98) and between NDVI and CIre (r = 0.76). In contrast, correlations of SIF with the optical vegetation indices were rather weak, except for SIF760 and EVI (r = 0.60) and MSAVI (r = 0.61). The two SIF bands themselves were moderately correlated with each other (r = 0.58). LST was weakly correlated with most optical indices, including NDVI and EVI. NDWI was negatively correlated with most greenness indices and SIF, but positively correlated with basal area (r = 0.47), LST (r = 0.44), and PRI (r = 0.31). Basal area was positively correlated with LST (r = 0.43) but negatively correlated with most greenness indices and SIF (r between *↑*0.24 and *↑*0.43, S2, see also Fig. S5 and section 3.5 below). CCI was weakly correlated with the other canopy variables, except for a moderate positive relationship with PRI (r = 0.34) and moderate negative relationships with SIF760 (r = *↑*0.33) and CIre (r = *↑*0.35).

Separating the data by class revealed differences in the correlation structure between needle-leaved and broadleaved tree species (Fig. S2b,c). SIF–greenness relationships were stronger in broadleaved than in needle-leaved tree species, while the latter showed stronger associations among NDWI, CIre, and PRI. Several correlations observed in the pooled dataset weakened when analyses were conducted within classes, particularly those involving basal area. This indicates that some pooled associations reflected differences between rather than within tree classes in the two variables.

In the Bechstedt experiment, across all functional groups, we observed strong positive correlations among vegetation greenness indices, indicating a consistent shared signal among these structural vegetation indices. However, their relationships with other remotely sensed canopy variables varied strongly among functional groups (Fig. S3). In small broadleaved tree species (Fig. S3b), greenness indices were positively correlated with CCI (NDVI–CCI: r = 0.75) and negatively with PRI (EVI–PRI: r = *↑*0.43; MSAVI–PRI: r = *↑*0.41). Early successional broadleaved tree species (Fig. S3c) had a distinct correlation structure with negative associations between basal area and thermal and greenness-related variables (ba–LST: r = *↑*0.48; ba–EVI: r = *↑*0.48; ba–MSAVI: r = *↑*0.46) and strong positive associations among pigment-related indices, including NDVI–EVI (r = 0.62), NDVI–CCI (r = 0.64), and CIre–PRI (r = 0.65). Late successional broadleaved tree species showed generally stronger coupling among canopy traits, with positive correlations between NDVI, EVI, MSAVI, and CCI (r = 0.45–0.70), as well as between CIre and PRI (r = 0.69). Needle-leaved tree species displayed the strongest correlations between greenness and chlorophyll-related indices, including NDVI–CCI (r = 0.82), NDVI–CIre (r = 0.75), and EVI/MSAVI–CCI (r = 0.71). Correlation patterns of Bechstedt needle-leaved tree species were similar to those observed in broadleaved tree species, with strong coherence among vegetation indices (Fig. S3b,c).

### 3.4 Multivariate structure of canopy variables

Next, we used PCAs to disentangle the differences across classes and functional groups further. In the Kaltenborn experiment, the first two principal components explained 44.7% and 17.0% of the total variance, respectively, accounting for 61.8% of the variation in the ten variables (Fig. 6a). PC1 was primarily associated with negative contributions from the greenness-related indices MSAVI, EVI and NDVI, as well as SIF760 and CIre, contrasting with positive contributions from NDWI and LST. PC2 was characterized mainly by negative contributions from PRI, NDVI and CCI, with LST contributing positively. The PCA scores showed a pronounced separation between needle-leaved and broadleaved tree species along PC1, with relatively limited overlap between the two classes.

**Fig. 6.**
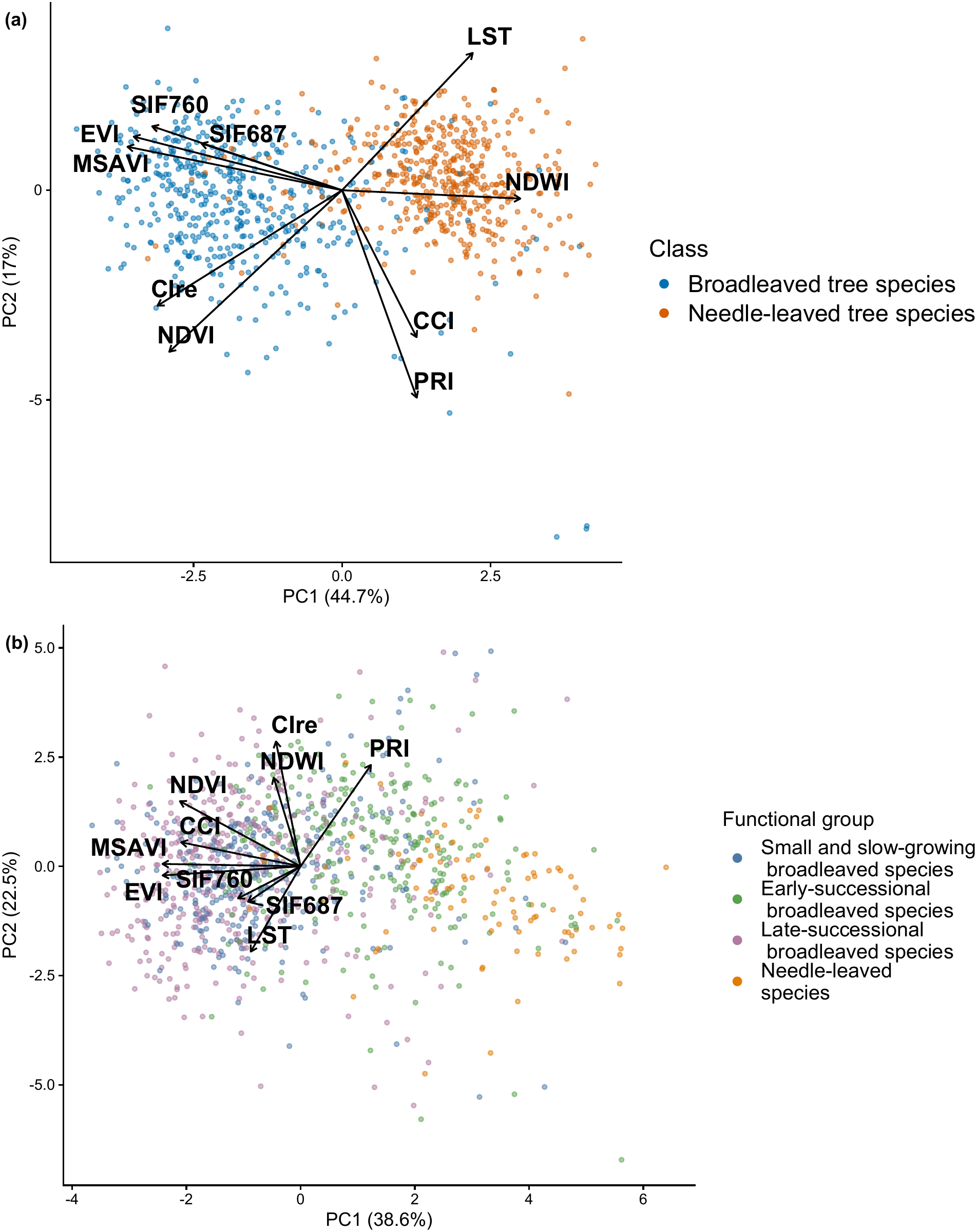
Principal component analysis (PCA) of the ten remote sensing-derived canopy variables. Basal area was excluded due to missing data and because it is a ground-based measurement. Variables were standardized prior to analysis. a) Kaltenborn. b) Bechstedt.

In the Bechstedt experiment, the first two principal components explained 38.6% and 22.5% of the total variance, respectively, accounting for 61.0% of the variation (Fig. 6b). PC1 was primarily associated with variation in the greenness-related indices EVI, MSAVI, NDVI and CCI, with both SIF variables and LST also contributing in the same direction. PC2 was characterized mainly by positive contributions from CIre, PRI, NDWI and NDVI, contrasting with negative contributions from LST and both SIF variables. The functional groups overlapped in PCA space, although needle-leaved species were generally separated from the broadleaved groups along PC1. The three broadleaved functional groups overlapped considerably, however, with a tendency to separate along the first PC (Fig. 6b). When sub-setting the data and only including the broad-leaved tree species, it became more clearly visible that the early-successional tree species separated from the other two functional groups along PC1 (Fig. S4), which was associated with the greenness-related vegetation indices EVI, MSAVI, NDVI and CCI.

Overall, the PCA revealed broadly similar correlation structures among the remotely sensed canopy variables in the two experiments (Fig. 6), with greenness-related indices contributing strongly to the first principal component at both sites. However, the distribution of observations differed between experiments. At Kaltenborn, the PCA scores showed a pronounced separation between broadleaved and needle-leaved species along PC1, whereas the four functional groups at Bechstedt showed substantial overlap, despite a tendency for needle-leaved species to have higher PC1 values.

### 3.5 Relationships between cumulative basal area and greenness-related indices as proxies for forest productivity

Finally, we investigated the relationship of greenness-related vegetation indices with cumulative basal area. Building on the correlation analyses presented in Section 2.3, we estimated site-specific slopes from linear models fitted across all classes, functional groups, species and diversity levels (Fig.S5). NDVI, EVI and MSAVI were significantly negatively associated with cumulative basal area at both sites (all Benjamini–Hochberg-adjusted *P <* 0.001), with consistently stronger negative slopes in Bechstedt than in Kaltenborn (NDVI: *↑*0.044 vs. *↑*0.167; EVI: *↑*0.424 vs. *↑*0.968; MSAVI: *↑*0.409 vs. *↑*0.879). CIre also declined significantly with increasing basal area in Kaltenborn (slope = *↑*5.13, adjusted *P <* 0.001), whereas no significant relationship was detected in Bechstedt (slope = 0.016, adjusted *P <* 0.979,; Fig.S5).

To test whether the overall relationships between cumulative basal area and greenness-related indices were consistent across individual species, we conducted separate linear regressions for each species within each experiment (Fig. S6). All regressions were restricted to patches for which basal area measurements were available. The species-specific analyses revealed more heterogeneous patterns than the pooled analyses, with negative relationships not consistently observed across all species (Fig. S6). In Kaltenborn, the only significant species-specific relationship was a negative association between NDVI and basal area for *P. menziesii* (slope = *↑*0.022, *P* = 0.0128). No significant relationships were detected for EVI, MSAVI or CIre in any of the four species. In Bechstedt, NDVI showed significant negative relationships with basal area for *B. pendula* (slope = *↑*0.088, *P* = 0.0196) and *P. sylvestris* (slope = *↑*0.199, *P* = 0.0142). EVI and MSAVI showed significant negative relationships for the same four species: *B. pendula* (EVI: slope = *↑*0.682, *P <* 0.001; MSAVI: slope = *↑*0.665, *P <* 0.001), *F. excelsior* (EVI: slope = *↑*1.21, *P <* 0.001; MSAVI: slope = *↑*1.30, *P <* 0.001, *P. sylvestris* (EVI: slope = *↑*0.714, *P* = 0.0034; MSAVI: slope = *↑*0.745, *P* = 0.0026) and *P. avium* (EVI: slope = *↑*0.685, *P <* 0.001; MSAVI: slope = *↑*0.616, *P <* 0.001). For CIre, no significant species-specific relationships were detected in Kaltenborn, whereas in Bechstedt, a significant positive relationship was found only for *T. cordata* (slope = 13.0, *P* = 0.0084).

Overall, increasing cumulative basal area was not consistently accompanied by increasing canopy greenness across either experiment. Because we fit many species-specific regressions, we are cautious in interpreting these individual p-values; but it is clear that greenness indices in the present experiments are opposite to the positive association assumed in studies suggesting the use of NDVI as a productivity measure.

## 4 Discussion

### 4.1 Weak effects of tree species diversity on community-level canopy variables

Tree species richness generally increases community-level functioning in forest biodiversity experiments and often also in comparative studies in natural forests (seeLiu et al. (2026) for a comprehensive recent review). Some previous studies using airborne canopy measurements in forest biodiversity experiments did find positive relationships for example between tree species richness and canopy structural complexity (Ray et al., 2024, 2023; Deng et al., 2025). It has also been demonstrated that remotely sensed spectral information can quantify net biodiversity effects on stem biomass and canopy nitrogen (Williams et al., 2020). We therefore hypothesized that tree species diversity would alter remotely sensed canopy signals at community level, but found little evidence for it. In the Kaltenborn experiment none of the tested relationships was significant (see Fig. 2) and in the Bechstedt experiment only the two canopy variables NDWI and PRI were positively affected at community level by functional diversity (see Fig. 3).

One possible reason for the weak effects of tree species diversity on community-level canopy variables is that we used a patch-wise planting scheme, where patches of monoculture tree stands were mixed to obtain plot-level mixed-species tree stands. Under this planting scheme, community-level effects of species richness or functional diversity may take longer to develop than in experiments where species are mixed at the individual level, such as in Huang et al. (2018). However, tree species richness had significant effects on most canopy variables in the Kaltenborn experiment at the species level; and similar clearly significant effects at functional group level were observed in the Bechstedt experiment (see discussion below). Another explanation is the lower statistical power of community-level analyses, which included only 16 or 25 plots, compared with 958 and 1100 patches in the species-level analyses at Kaltenborn and Bechstedt, respectively. Most importantly, weak community-level responses likely reflect contrasting responses among species and functional groups within communities.

### 4.2 Species differences and effects of tree species diversity on species-level canopy variables

Differences between tree species in remotely sensed canopy variables have often been observed, in particular if species are also phylogenetically widely separated (Schweiger et al., 2018; Meireles et al., 2020). Such differences are the basis for detecting broader taxonomic groups or species by satellite or airborne imaging (van Moorsel et al., 2026; Fassnacht et al., 2016). However, van Moorsel et al. (2026) showed that individual species can express significant variation in spectral features in response to tree species diversity. We therefore also hypothesized that species identity together with tree species richness or functional diversity would alter the canopy variables derived from spectral features or from other remotely-sensed signals analyzed here at species level. Species-specific responses to tree species diversity might than also explain the absence of canopy-level responses discussed above.

Across the canopy variables, species identity explained substantially more variation than tree species richness. Most vegetation indices clearly separated broadleaved tree species from needle-leaved tree species, and additional contrasts among species were also highly significant. For example, broadleaved tree species consistently had higher NDVI, EVI, and CIre values than needle-leaved tree species, reflecting differences in leaf morphology, leaf biochemistry, canopy architecture and thus optical properties between broadleaved and needle-leaved species (Meireles et al., 2020; Croft et al., 2014; van Moorsel et al., 2026). That these differences were so pronounced across multiple indices suggests that canopy spectral properties in this system are primarily determined by species-specific physiological and structural traits. This is consistent with previous work demonstrating functional identity to be the primary driver of biodiversity effects on productivity in young tree communities (Grossiord et al., 2013; Tobner et al., 2016).

Species responses to diversity can either increase or reduce interspecific differences. At Kaltenborn, differences between needle-leaved and broadleaved species increased with richness for NDWI, NDVI and CIre, whereas functional-group differences generally decreased with functional diversity at Bechstedt. Such patterns indicate differential trait plasticity of species growing in mixtures, where species adjust their physiological or structural characteristics when growing alongside other species (Yang et al., 2026; Niklaus et al., 2017). Increasing divergence would be expected when it increases niche differences between species, whereas increasing similarity would be expected when it decreases fitness differences between species (Liu et al., 2026; Chen et al., 2025). Such responses may differ depending on the phylogenetic level considered (Allan et al., 2013). In our study this was reflected by the contrasting responses of CCI and CIre, with increased differences between needle-leaved tree species and broadleaved tree species, but reduced differences between species within these two groups in mixtures in the Kaltenborn experiment (see Fig. 2I-J). Increasing divergence in remotely sensed canopy variables with richness, potentially leading to increasing canopy structural differentiation and complementarity among species in mixed stands, is consistent with findings in previous biodiversity experiments (Deng et al., 2025). The increasing similarity with increasing functional diversity in the Bechstedt experiment may reflect stronger environmental filtering among a larger number of species and an already high functional diversity set by the design. Although spectral indices are only proxies for physiological traits, these patterns suggest that interspecific interactions alter trait expression without necessarily affecting community-level functioning.

### 4.3 Species identity alters the relationship among canopy variables

Relationships among remotely sensed variables differed between tree species from different phylogenetic or functional groups (Fig. S2, Fig. S3). Greenness indices such as NDVI, EVI, and MSAVI were consistently correlated across groups, likely because they all capture aspects of canopy greenness. In contrast, their relationships with physiological variables, including SIF, PRI, and canopy temperature, differed substantially between functional groups. These physiological indicators may be influenced by species-specific regulation of photosynthesis, pigment dynamics, and water use, although they may also be affected by spectral signatures of the structural differences between needle-leaved and broadleaved trees. However, because both SIF and LST data were acquired at the same time with a comparable spatial resolution, contrasting structural interferences affecting the retrieval of SIF and LST are unlikely. For example, at Kaltenborn, needle-leaved tree species showed a negative and broadleaved tree species a positive correlation between SIF and LST, with similar but weaker results for Bechstedt (Fig. S2, Fig. S3). While a negative relationship is consistent with previous literature demonstrating that thermal stress reduces fluorescence yield and photosynthetic efficiency by altering energy partitioning within the photosystems (Paul-Limoges et al., 2018), the positive relationship for broadleaved tree species in both experiments is unexpected. An airborne experiment (Damm et al., 2022) demonstrated that SIF can show a double response to stress with an initial physiologically-caused increase due to downregulation of photosynthesis (also predicted by a leaf-level model: van der Tol et al. (2009)), followed by a decline due to degrading chlorophyll pigments. This would suggest that different manifestations of stress were detected for coniferous (established stress) and broadleaved (immediate stress) species. Another possible explanation is that canopy temperatures remained below the threshold at which thermal stress substantially constrains photosynthesis. At the temperatures observed in our experiments during data acquisition (around 30° C), higher leaf temperatures may instead have coincided with conditions that favoured photosynthetic activity, while stronger thermal stress and stomatal closure may only occur at higher temperatures. The positive SIF–temperature relationship may reflect the absence of severe heat stress rather than a more general positive effect of temperature on photosynthesis. The needle-leaved tree species *P. abies* exhibited the highest canopy temperatures and lowest SIF, consistent with its adaptation to cooler climates and reduced physiological performance under elevated temperatures (Orešković et al., 2024). The needle-leaved tree species *P. menziesii* showed similarly warm canopies and low SIF (Fig. 4). In contrast, the broadleaved tree species generally maintained cooler canopies (particularly *Q. petraea*, whose canopy temperature decreased with increasing species richness) so that the positive relationship of canopy temperature with SIF may indicate increased photosynthetic performance across temperature ranges not reaching detrimentally high levels. These contrasting responses suggest that species identity influences both canopy temperature and how canopy thermal conditions translate into photosynthetic functioning. It has been reported recently that differences in canopy architecture, including crown structure and the distribution of sunlit and shaded foliage, can influence observed thermal patterns (Adams et al., 2025). Greater diversity can increase canopy structural complexity (Williams et al., 2017; Jucker et al., 2015) and alter the distribution of sunlit and shaded foliage, thereby modifying the canopy energy balance and remotely sensed surface temperature.

The contrasting relationships between SIF and other canopy variables may also reflect differences in canopy structure and light distribution between tree species. This interpretation is supported by previous leaf-level measurements in the Kaltenborn experiment, where chlorophyll fluorescence responses to tree diversity differed among species, suggesting that community context modifies photosynthetic functioning at both leaf and canopy scales (Pollastrini et al., 2014). Although SIF is closely linked to photosynthetic activity (Pickering et al., 2022), the relationship between SIF and GPP is influenced by the differently responding yields of SIF and photosynthesis, but also by canopy structural properties, including leaf area distribution and vegetation structure (Shi et al., 2025; Forrester et al., 2019). Therefore, variation in canopy architecture between needle-leaved tree species and broadleaved tree species may also have contributed to the observed differences in SIF relationships with greenness indices.

At Kaltenborn, several correlations between remotely-sensed variables apparent in the whole dataset weakened after separating needle-leaved tree species from broadleaved tree species (Fig. S2). This indicates that correlations between canopy traits can emerge primarily from differences among rather than within functional groups. Consequently, analyses that ignore forest composition may overestimate the generality of relationships between remotely-sensed traits and ecosystem functioning. This also suggests that empirical relationships among remotely-sensed variables are not necessarily transferable across forest types. Models calibrated using one functional composition may perform poorly in communities with contrasting species identities because physiological relationships among spectral traits differ systematically among functional groups.

### 4.4 Remotely-sensed greenness indices are not direct proxies for forest productivity

Vegetation indices such as NDVI and EVI are widely used to characterize canopy greenness, vegetation activity, and structural properties of ecosystems and are often used as proxies of productivity-related variables such as GPP or biomass accumulation (Phillips et al., 2008; Birky, 2001; Zhang et al., 2022). Cumulative basal area integrates tree growth over time and is Here, we tested whether remotely-sensed canopy traits were consistently related to cumulative basal area as an indicator of forest productivity (Grossiord et al., 2013; Hessburg et al., 2016), and whether these relationships differed among tree species belonging to different phylogenetic or functional groups.

Contrary to expectations, cumulative basal area was negatively instead of positively correlated with spectral indices commonly used to remotely assess canopy productivity. This indicates that under experimental conditions — where growth variation is primarily due to species differences and differences in designed tree diversity rather than to abiotic environmental heterogeneity — greater accumulated tree growth can be negatively associated with canopy greenness. Remotely sensed vegetation indices and ground-based measures of tree growth thus capture different aspects of ecosystem functioning. While spectral indices primarily provide a snapshot of current canopy variables, including leaf area, pigment content, nutrient composition (especially nitrogen), and canopy structure, cumulative basal area represents an integrated structural outcome of tree growth over years. Although positive relationships between NDVI and tree growth have been reported from observational studies, these relationships vary substantially with temporal scale, seasonality, forest type, and environmental conditions (le Maire et al., 2011; Vicente-Serrano et al., 2016). Correlations between a vegetation index measured at a single point in time and cumulative timber yield should be interpreted cautiously, as they may reflect indirect relationships mediated by stand age, structure, species composition, or growing conditions rather than a direct relationship between canopy greenness and accumulated woody biomass. Thus, without complete time series of basal area and canopy greenness, it is difficult to determine when, and whether, differences in greenness indices reflect differences in biomass allocation among species.

NDVI is among the most widely used vegetation indices (Pettorelli et al., 2005). However, NDVI is prone to saturation at high levels of canopy density and leaf area index (LAI), limiting its ability to capture variation among dense forest canopies (Huete et al., 2002). Once canopy cover approaches saturation, further increases in biomass or productivity may result in no further, or only minor, changes in NDVI values (Huete et al., 2002). This limitation is particularly relevant for mature forests, where structural differences among species or stands may not be fully represented by greenness-based indices. The enhanced vegetation index (EVI) was developed to improve sensitivity in high-biomass ecosystems by reducing atmospheric and background effects and by partially mitigating saturation effects observed in NDVI (Huete et al., 2002). Nevertheless, EVI remains a greenness-based index rather than a direct measurement of productivity, and its relationship with forest functioning can still depend on canopy structure, species composition, and environmental conditions (Koch et al., 2025). Methodological aspects can additionally confound the sensitivity of these indices to temporally accumulated productivity. Large growth reflected by high basal area can be manifested in more heterogeneous canopies in terms of crown height and shape, causing different shadow characteristics that, in turn, can affect the retrieval of greenness indices (Damm et al., 2015b; Fawcett et al., 2018). Thus, vegetation indices may be less suitable as universal, species-independent indicators of canopy functioning than their widespread use might suggest.

### 4.5 Limitations and conclusions

There are several limitations to our study. First, spectral indices and SIF integrate multiple structural and physiological processes rather than representing single plant functions (Barton and North, 2001; Gamon et al., 1995). Here, we show a case in which greenness indices often do not correlate positively to a more direct measure of productivity, but we do not know exactly what drives observed differences in index values in our dataset. Second, the relatively young age may limit the transferability of our findings to mature forests. However, the unique design of the BIOTREE experiments at Kaltenborn and Bechstedt ensures that canopy measurements can be assigned to particular species at patch level, avoiding species-identification issues within mixed pixels, as well as confounding effects of abiotic environmental heterogeneity affecting observational studies in natural or managed forests. Only with such experimental studies is it possible to resolve variation among tree species and identify the mechanisms underlying species-specific responses within mixtures at plot scales. Expanding airborne and satellite observations across further biodiversity experiments (Veryard et al., 2023; Deng et al., 2025) or provenance trials (Csilléry et al., 2026) where remotely-sensed variables can be clearly assigned to single species, would therefore help evaluate the generality of these relationships across ecosystems and climatic conditions. Future studies combining remote sensing with physiological measurements and detailed canopy structure will be needed to better understand the mechanisms driving variation in spectral signals.

Our results demonstrate that species identity influences remotely-sensed canopy traits as measured by commonly used indices as well as SIF, and their correlations, and that traits of different species, so measured, significantly vary in their responses to plot-level tree species richness or functional diversity. These differences between tree species belonging to different phylogenetic or functional groups highlight the importance of considering species composition when interpreting remotely sensed indicators of forest functioning. Forest biodiversity experiments provide valuable platforms for linking remote-sensing approaches with biodiversity–ecosystem functioning research.

## Author contributions

**Sofia J. van Moorsel**: conceptualization, formal analysis, investigation, methodology, visualization, project administration, writing – original draft, writing – review and editing. **Bernhard Schmid**: conceptualization, formal analysis, investigation, methodology, writing - original draft, writing - review and editing **Michael Niederberger**: data curation, methodology, formal analysis, visualization, writing – review and editing **Michael Scherer-Lorenzen**: methodology, writing – review and editing **Tiziana L. Koch**: writing – review and editing **Daniel Kopkane**: methodology, writing – review and editing **Uwe Rascher**: methodology, writing – review and editing **Alexander Damm**: conceptualization, methodology, funding acquisition, writing – review and editing **Meredith C. Schuman**: conceptualization, funding acquisition, writing – review and editing.

## Acknowledgements

This research was funded by the NOMIS foundation grant “Remotely sensing ecological genomics” to M. E. Schaepman (M. C. Schuman is co-investigator) and by the Swiss National Science Foundation (SNSF), Switzerland, project Fluo4Eco (grant number 197243). The BIOTREE experiment has been established by the Max-Planck-Institute for Biogeochemistry Jena, Germany, and we are very grateful to Prof. Dr. Ernst-Detlef Schulze for making this project possible. The authors thank Dietrich Mackensen and Klaus Hahner (Federal Forestry Office Thüringer Wald—Bundesforstamt Thüringer Wald, Bad Salzungen) for support and site maintenance.

## Data availability statement

Data used in this study will be made publicly available on Dryad upon acceptance of the manuscript [link and DOI tbd]. Code will be available on a code repository (Zenodo, link and DOI tbd).

## Supporting Information

Supporting information can be found online.

