## Supporting Information for "Effects of tree species diversity and identity on remotely sensed canopy variables in two forest biodiversity experiments"

**Table of contents**

**Table S1.** ANOVA of effects of tree diversity and class on canopy variables and basal area at Kaltenborn.
**Table S2.** ANOVA of effects of functional diversity and plant functional group on canopy variables and basal area at Bechstedt.
**Figure S1.** Effects of species richness and functional diversity on the ten canopy variables and basal area.
**Figure S2.** Correlation structure of canopy variables and basal area in the Kaltenborn experiment.
**Figure S3.** Correlation structure of canopy variables in the Bechstedt experiment.
**Figure S4.** PCA für angiosperms in Bechstedt.
**Figure S5.** Relationships between cumulative basal area and remotely sensed spectral indices for greenness across the two study sites.
**Figure S6.** Relationships between cumulative basal area and remotely sensed vegetation indices across tree species.

**Table S1.** Effects of species richness, class and species identity within class on canopy variables and basal area at the Kaltenborn site where species richness was manipulated. Results from linear mixed-effects models with each canopy variable as the response. Models included log-transformed species richness (logsr), class (CL), contrasts (PI for *P. abies* vs *P. menziesii* and FA for *F. sylvatica* vs. *Q. petraea*), and richness × identity interactions as fixed effects, with plot and subplot included as nested random effects. Values shown are *F* statistics and *P*-values from model analysis of variance (ANOVA) with P values $< 0.05$ in **bold**. N = 16 plots and 32 subplots; ba: basal area in $m^2$ $ha^{-1}$, LST: Land Surface Temperature in K; indices are given in Table 1.

| Response | *F* logsr | *F* CL | *F* PI | *F* FA | *F* logsr:CL | *F* logsr:PI | *F* logsr:FA | *P* logsr | *P* CL | *P* PI | *P* FA | *P* logsr:CL | *P* logsr:PI | *P* lo |
|---|---|---|---|---|---|---|---|---|---|---|---|---|---|---|
| Basal area | 0.8 | 258.18 | 7.56 | 7.73 | 17.29 | 1.98 | 2.97 | 0.388 | <**0.001** | **0.006** | **0.006** | <**0.001** | 0.160 | |
| LST | 1.23 | 331.28 | 4.18 | 1.91 | 5.27 | 4.09 | 2.36 | 0.286 | <**0.001** | **0.041** | 0.168 | **0.022** | **0.043** | |
| NDWI | 0 | 850.16 | 169.43 | 3.66 | 9.65 | 0.88 | 1.72 | 0.968 | <**0.001** | <**0.001** | 0.056 | **0.002** | 0.35 | |
| NDVI | 1.22 | 497.75 | 4.91 | 0.42 | 13.6 | 0.05 | 0.38 | 0.288 | <**0.001** | **0.027** | 0.516 | **0.001** | 0.828 | |
| EVI | 1.19 | 156.42 | 9.3 | 2.73 | 1.66 | 1.04 | 0.44 | 0.294 | <**0.001** | **0.002** | 0.099 | 0.198 | 0.934 | |
| MSAVI | 0.62 | 190.55 | 3.26 | 2.53 | 5.36 | 0.56 | 0.31 | 0.444 | <**0.001** | **0.044** | 0.112 | **0.021** | 0.675 | |
| SIF760 | 0.28 | 84.46 | 0.36 | 2.55 | 2.88 | 0.53 | 2.21 | 0.606 | <**0.001** | 0.551 | 0.11 | **0.021** | 0.615 | |
| SIF687 | 0.57 | 34.79 | 10.12 | 0.22 | 5.34 | 0.16 | 0.68 | 0.464 | <**0.001** | **0.002** | 0.642 | **0.021** | 0.688 | |
| CCI | 0.17 | 1.15 | 92.5 | 184.52 | 2.89 | 22.16 | 0.68 | 0.683 | 0.284 | <**0.001** | <**0.001** | 0.09 | **0.001** | |
| CIre | 0.07 | 469.93 | 36.66 | 10.72 | 10.9 | 10.93 | 4.29 | 0.79 | <**0.001** | <**0.001** | **0.001** | <**0.001** | <**0.001** | |
| PRI | 0 | 15.15 | 41.24 | 77.03 | 1.13 | 0.48 | 0.8 | 0.968 | **0.001** | <**0.001** | <**0.001** | 0.287 | 0.488 | |

**Table S2.** Effects of functional diversity and plant functional group on canopy variables and basal area at the Bechstedt site where functional diversity was manipulated at a constant richness of four species per plot. Results from linear mixed-effects models fitted separately for each canopy response variable. Models included functional diversity (FD), plant functional group (FG), and their interaction (FD × FG) as fixed effects, with plot identity (PL) included as a random effect. Functional groups comprised small broadleaved tree species, early-successional broadleaved tree species, late-successional broadleaved tree species, and needle-leaved tree species. Values shown are *F* statistics and corresponding *P*-values from model ANOVA with P values $< 0.05$ in **bold**; ba: basal area in $m^2$ $ha^{-1}$, LST: Land Surface Temperature in K; indices are given in Table 1.

| Response | *F* FD | *F* FG | *F* FD:FG | *P* FD | *P* FG | *P* FD:FG |
|---|---|---|---|---|---|---|
| Basal area | 2.00 | 99.30 | 3.72 | 0.171 | <**0.001** | **0.011** |
| LST | 2.25 | 9.13 | 1.78 | 0.148 | <**0.001** | 0.149 |
| NDWI | 5.63 | 3.92 | 4.73 | **0.026** | **0.008** | **0.003** |
| NDVI | 0.01 | 189.51 | 13.27 | 0.908 | <**0.001** | <**0.001** |
| EVI | 1.50 | 133.08 | 9.74 | 0.232 | <**0.001** | <**0.001** |
| MSAVI | 1.30 | 160.14 | 13.69 | 0.265 | <**0.001** | <**0.001** |
| SIF760 | 2.01 | 7.22 | 3.42 | 0.169 | <**0.001** | **0.006** |
| SIF687 | 2.34 | 3.05 | 6.51 | 0.140 | **0.028** | **0.001** |
| CCI | 0.28 | 150.44 | 10.49 | 0.602 | <**0.001** | <**0.001** |
| CIre | 2.75 | 24.23 | 5.99 | 0.111 | <**0.001** | <**0.001** |
| PRI | 5.82 | 29.04 | 5.83 | **0.024** | <**0.001** | <**0.001** |

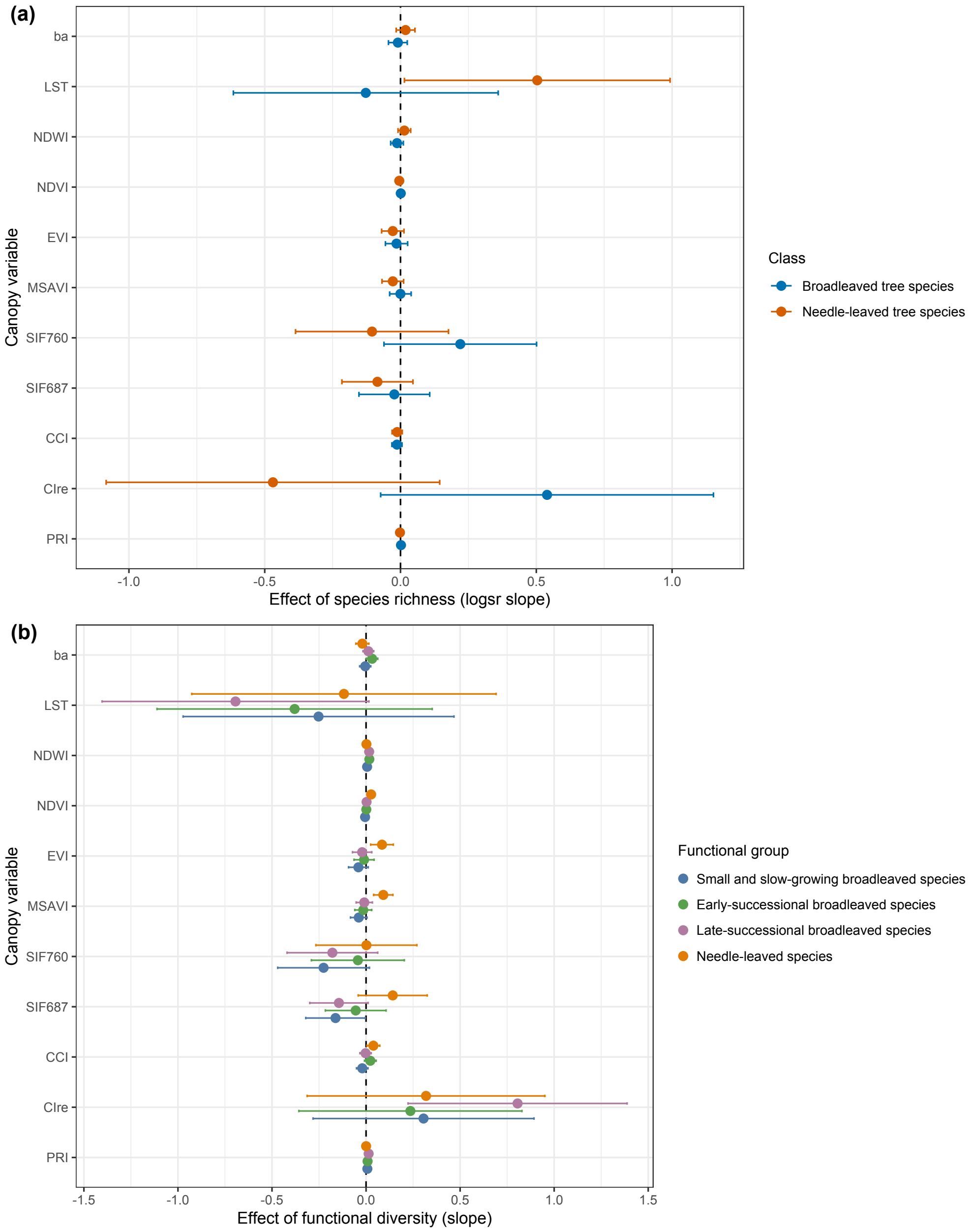


**Fig. S1.** Effects of species richness and functional diversity on the ten canopy variables and basal area (ba). (a) Kaltenborn experiment: Estimated slopes (estimated marginal trends using emtrends function in the emmeans package) of plot-level species richness (log-transformed species richness) for canopy variables measured for broadleaved tree species and needle-leaved tree species. (b) Bechstedt experiment: Estimated slopes of plot-level functional diversity on canopy variables and basal area (ba) are shown separately for the four functional groups: small broadleaved tree species, early-successional broadleaved tree species, late-successional broadleaved tree species, and needle-leaved tree species. Points show model-estimated slopes ± 95% confidence intervals. Slopes represent the relationship between plot-level species richness and canopy variables within each class or functional group; they do not represent independent species-level richness effects. Positive values indicate increasing canopy responses with increasing richness or functional diversity, whereas negative values indicate decreasing responses. Confidence intervals overlapping zero indicate effects that are not distinguishable from zero. ba: basal area in $m^2$ $ha^{-1}$, LST: Land Surface Temperature in K; indices are given in Table 1.

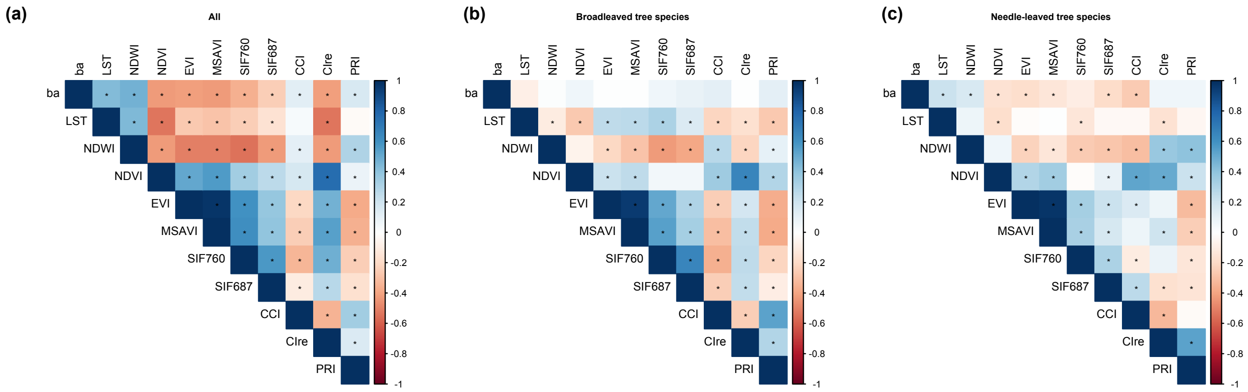


**Fig. S2.** Correlation structure of canopy variables and basal area in the Kaltenborn experiment. Pearson correlation matrices for (a) all species, (b) broadleaved tree species, and (c) needle-leaved tree species. Colours indicate the magnitude and direction of correlations, with significant relationships ($P < 0.05$) indicated with a star; ba: basal area in $m^2$ $ha^{-1}$, LST: Land Surface Temperature in K; indices are given in Table 1.

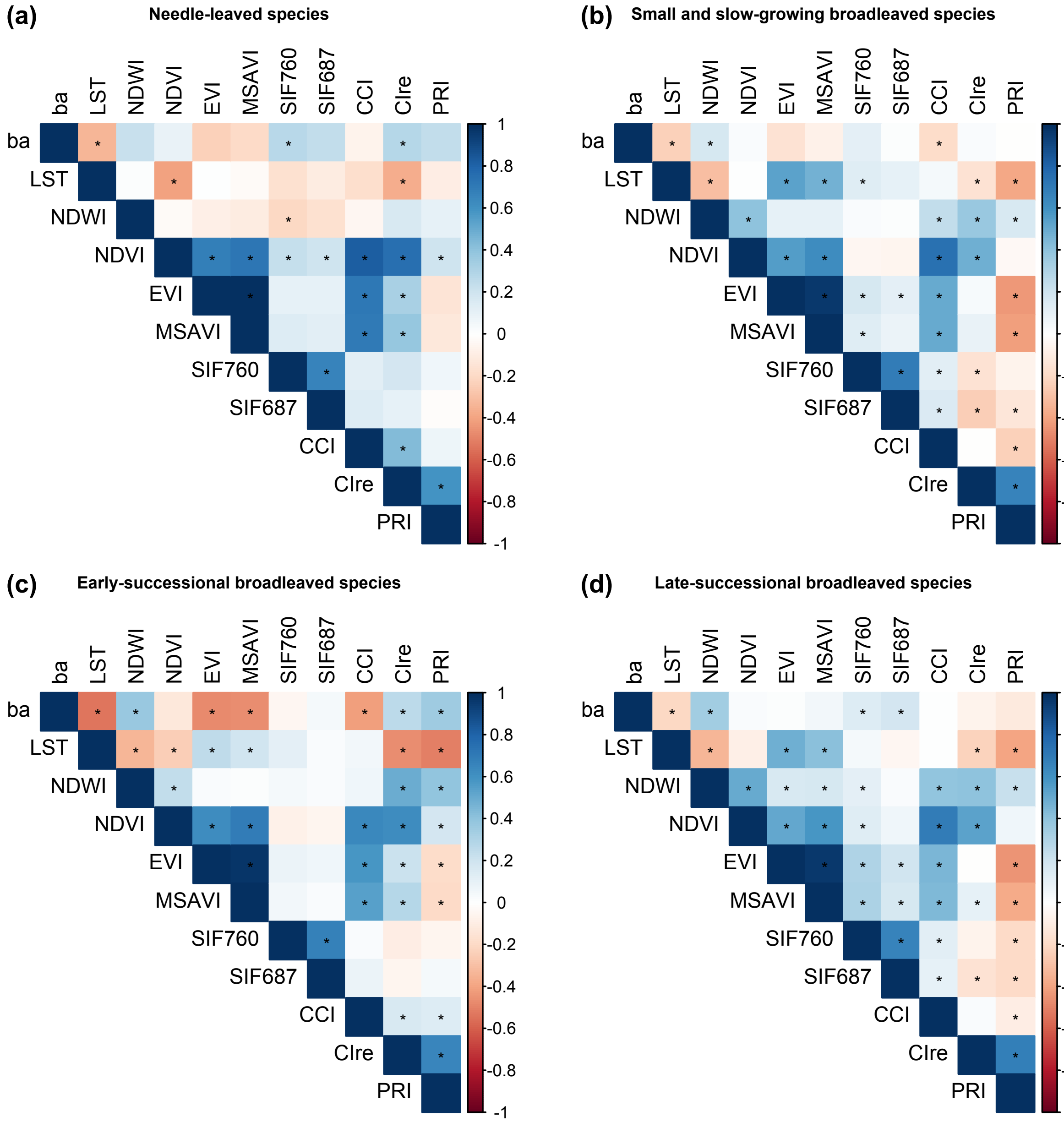


**Fig. S3.** Correlation structure of canopy variables in the Bechstedt experiment. Pearson correlation matrices of canopy variables and basal area for (A) needle-leaved tree species, (B) small broadleaved tree species, (C) early-successional broadleaved tree species and (D) late-successional broadleaved tree species. Colours indicate the magnitude and direction of correlations, with significant relationships ($P < 0.05$) indicated with a star; ba: basal area in $m^2$ $ha^{-1}$, LST: Land Surface Temperature in K; indices are given in Table 1.

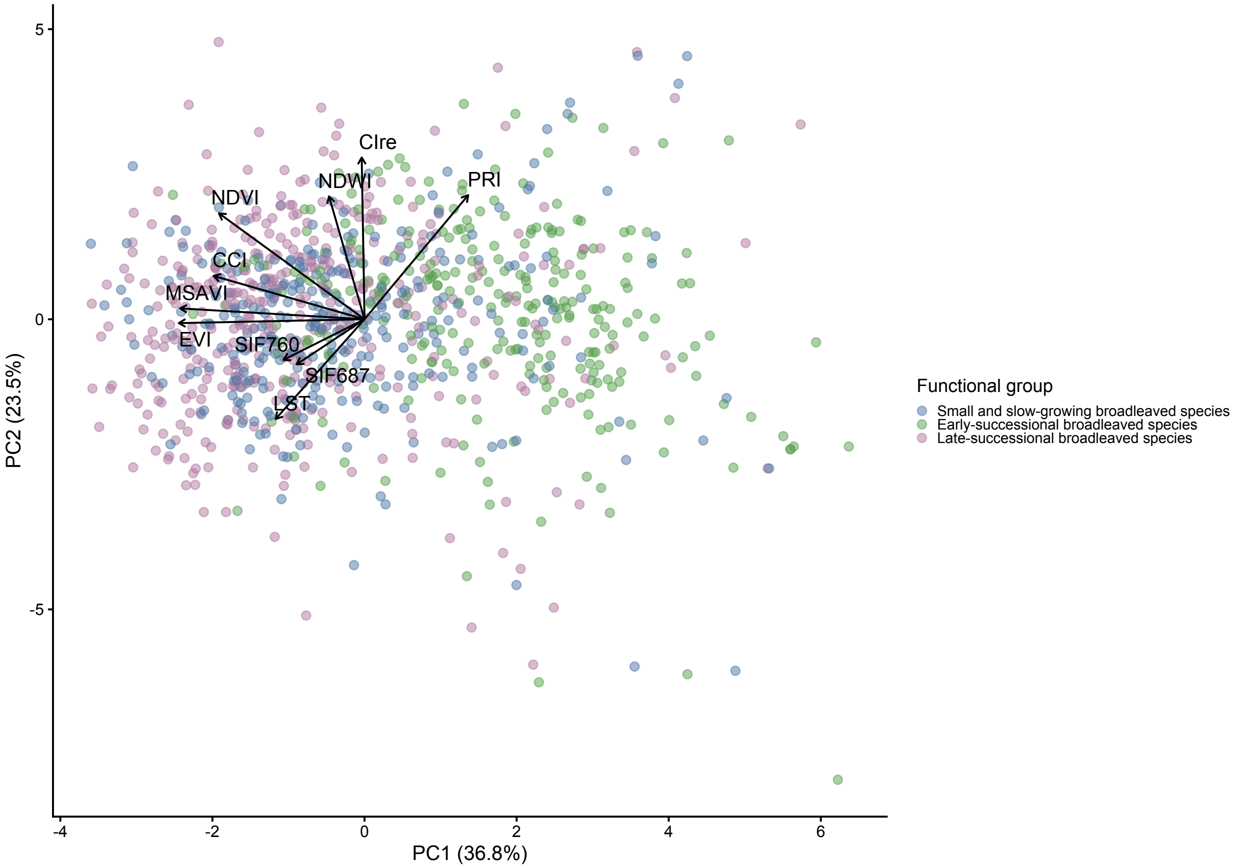


**Fig. S4.** PCA for the subset of the broad-leaved tree species in Bechstedt; LST: Land Surface Temperature in K; indices are given in Table 1.

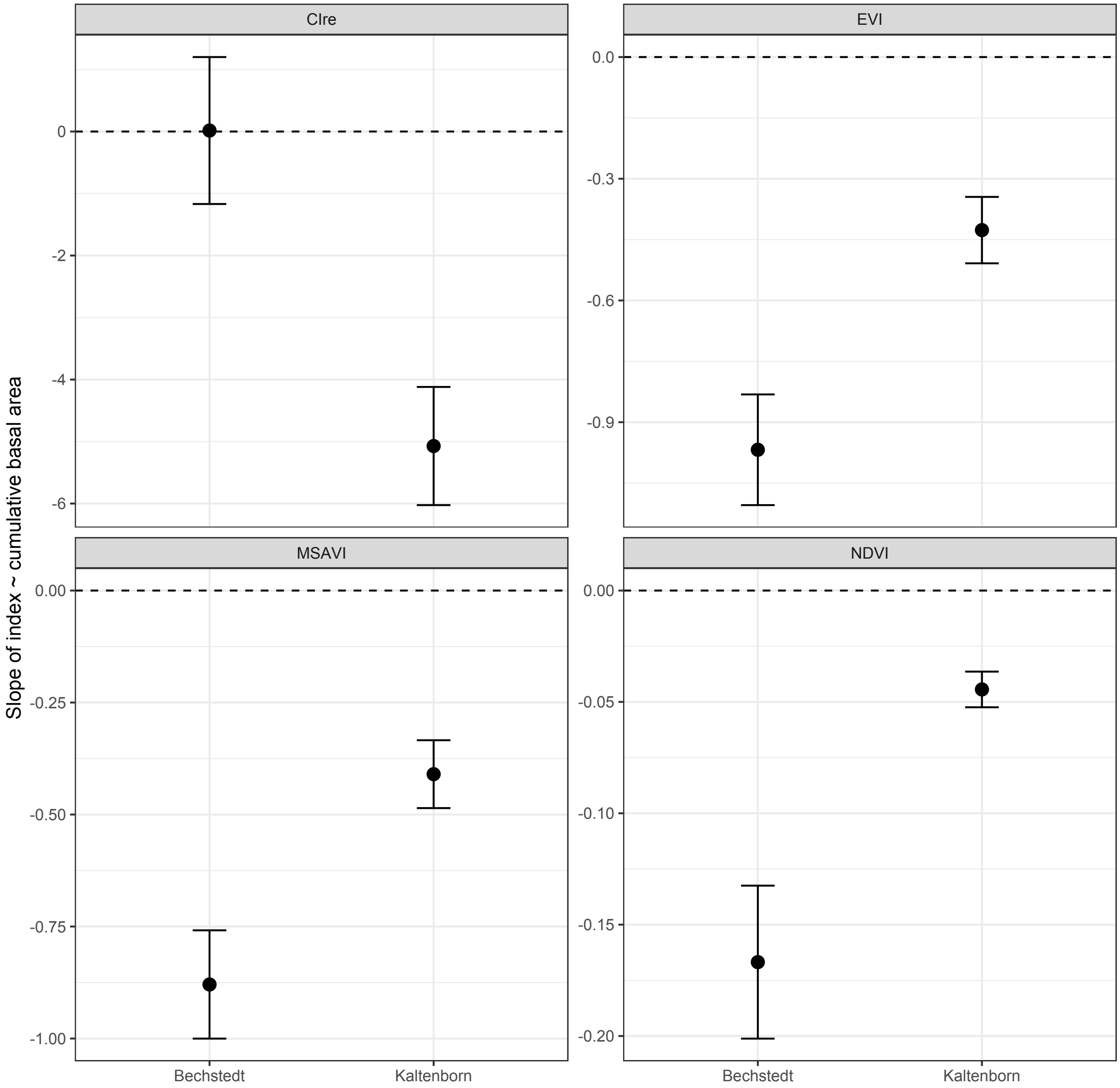


**Fig. S5.** Relationships between cumulative basal area in $m^2$ $ha^{-1}$ and remotely sensed spectral indices (CIre, EVI, NDVI and MSAVI) across the two study sites. Overall slopes were estimated from linear models relating each spectral index to cumulative basal area separately for Kaltenborn and Bechstedt. Points represent individual patch observations pooled across all species within each site. Error bars indicate 95% confidence intervals of the estimated slopes. Negative slopes indicate decreasing index values with increasing cumulative basal area. Indices are given in Table 1.

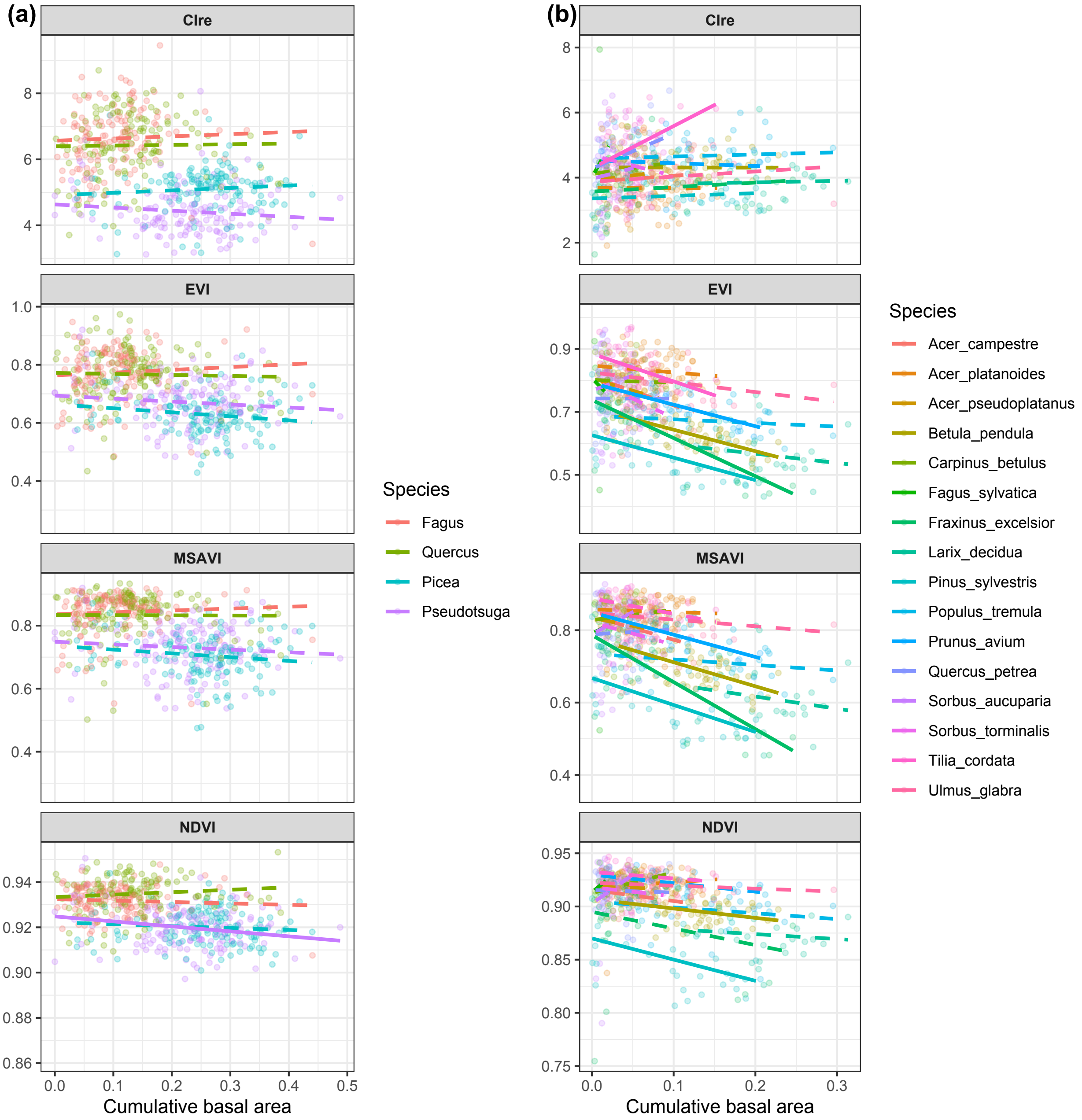


**Fig. S6.** Relationships between cumulative basal area in $m^2$ $ha^{-1}$ and remotely sensed vegetation indices (CIre, EVI, NDVI and MSAVI) for each tree species separately in (a) Kaltenborn and (b) Bechstedt. Points represent individual observations, coloured lines represent species-specific linear regressions. Solid lines indicate significant relationships ($P < 0.05$, non-adjusted), whereas dashed lines indicate non-significant relationships. This indicates whether each species-specific slope differs significantly from zero. It does not test whether slopes differ significantly between species. Indices are given in Table 1.
